# Cryo-EM structure of the ClpXP protein degradation machinery

**DOI:** 10.1101/638692

**Authors:** C Gatsogiannis, D Balogh, F Merino, SA Sieber, S Raunser

## Abstract

The ClpXP machinery is a two component protease complex performing targeted protein degradation in bacteria and eukaryotes. The complex consists of the AAA+ chaperone ClpX and the peptidase ClpP. The hexameric ClpX utilizes the energy of ATP binding and hydrolysis to engage, unfold and translocate substrates into the catalytic chamber of tetradecameric ClpP where they are degraded. Formation of the complex involves a symmetry mismatch, since hexameric AAA+ rings bind axially to the opposing stacked heptameric rings of the tetradecameric ClpP. Here we present the first high-resolution cryo-EM structure of ClpXP from *Listeria monocytogenes*. We unravel the heptamer-hexamer binding interface and provide novel insights into the ClpX-ClpP crosstalk and activation mechanism. The comparison with available crystal structures of ClpP and ClpX in different states allows us to understand important aspects of ClpXP’s complex mode of action and provides a structural framework for future pharmacological applications.

## Results and Discussion

Caseinolytic protease P (ClpP) represents a major proteolytic protein in prokaryotes and eukaryotes involved in protein homeostasis, bacterial pathogenesis as well as cancer progression (*1*–*3*). ClpP is highly conserved, essential for virulence and regulation of stress responses in several pathogenic bacteria and therefore considered as a promising therapeutic target for novel antibiotics (*4*). ClpP associates with diverse ATP-dependent AAA+ chaperones such as ClpX, ClpC and ClpA for the recognition, unfolding and digestion of substrate proteins (*5*). To date, a large fraction of research has been dedicated to functionally exploit ClpP and its cognate chaperones, foremost ClpX, in terms of their enzymatic activity, individual structures and conformational control.

Previous low resolution electron microscopy (EM) studies of ClpXP and ClpAP from *Escherichia coli* revealed that up to two hexameric ClpX chaperones bind to a ClpP tetradecameric barrel (*6*, *7*). Early on, the hexamer-heptamer interface fascinated researchers and several studies characterizing the role of putative interaction motifs have led to the proposition of models explaining the symmetry mismatch and functional interaction between the two proteins (*7*–*10*). Sequence alignments and mutational studies of AAA+ chaperones identified loops in ClpX, that interact with the hydrophobic clefts on the periphery of ClpP. They contain the highly conserved (I/L/V)-G-(F/L) motif and are essential for complex formation (*11*).

More recently, cyclic acyldespipeptides (ADEPs), a novel class of anti-bacterial compounds, have been identified to bind to the same peripheral hydrophobic clefts on ClpP and to induce the opening of the axial pores of ClpP (*4*, *12*–*14*). They stabilize ClpP in an “open” activated state in the absence of the chaperone, leading to unregulated proteolysis of substrates and finally to cell death. This suggests that the protruding loops in ClpX that contain the (I/L/V)-G-(F/L) motif, also called IGF loops, are sufficient to activate ClpP. It has also been speculated that this activation involves the opening of the axial pore to allow translocation of the substrate into the proteolytic chamber of ClpP (*15*).

Contacts between the pore-2 loops of ClpX and the N-termini of ClpP represent a second set of well-characterized interactions between ClpX and ClpP, which are, however, more dynamic and dependent on the nucleotide state of ClpX (*10*). A crucial function of the ClpP N-termini is to gate the entrance of the proteolytic chamber (*9*). Until now, the highly dynamic interaction between ClpX and ClpP, which is mediated by long flexible loops, posed a challenge to obtaining a high resolution structure. This limited our understanding of this important protein degradation machinery. Here we present the first high-resolution cryo-EM structure of ClpXP from *L. monocytogenes*.

### Cryo-EM structure of ClpXP1/2

In order to obtain a ClpXP complex that is suitable for structural studies, we used the heterotetradecameric ClpP1/2 complex from *Listeria monocytogenes.* Recent studies have revealed that this complex has a higher affinity to ClpX in comparison to the more conserved ClpP2 homocomplex (*16*, *17*) suggesting a superior stability of the heterooligomer. Since ClpP1/2 might cleave ClpX to some extent during sample preparation, we mutated one residue of the catalytic triad (S98A) in both ClpP isoforms to prevent cleavage. Furthermore, we mutated the nucleotide binding site of ClpX (E183Q) to allow ATP binding, but to prevent hydrolysis, which results in a tighter binding to ClpP (*18*, *19*).

We formed a complex of ClpX and ClpP1/2 and obtained a large fraction of ClpXP1/2 dimers (ClpP1/2-ClpX-ClpX-ClpP1/2) that were in equilibrium with ClpXP1/2 monomers (Supplementary Fig. 1a-c). It has been demonstrated before that two ClpX or ClpA hexamers can bind to one ClpP barrel from both sites, resulting in a ClpX-ClpP-ClpX or ClpA-ClpP-ClpA complex (*6*, *7*, *20*). However, ClpXP1/2 dimers (Supplementary Figure 1a-d) have, to our knowledge, not been described so far. We therefore concentrated our structural analysis first on these intriguing dimers and determined their structure by cryo-EM and single particle analysis using crYOLO (*21*) and SPHIRE (*22*) (Figure 1a-b, Supplementary Fig. 1e-g). Although the intrinsic flexibility of the complexes did not allow the determination of a high-resolution structure (Supplementary Video 1, Supplementary Figure 1e-g), the fitting of the crystal structure of ClpX into the cryo-EM density suggests that the flexible N-terminal zinc binding domains (ZBDs) of ClpX mediate the interaction between two ClpX hexamers (Figure 1c). While ZBD-deleted ClpX still associated with ClpP to a small extent, ClpX dimerization was completely abolished supporting our structural data (Supplementary Fig. 1a).

**Figure 1.**
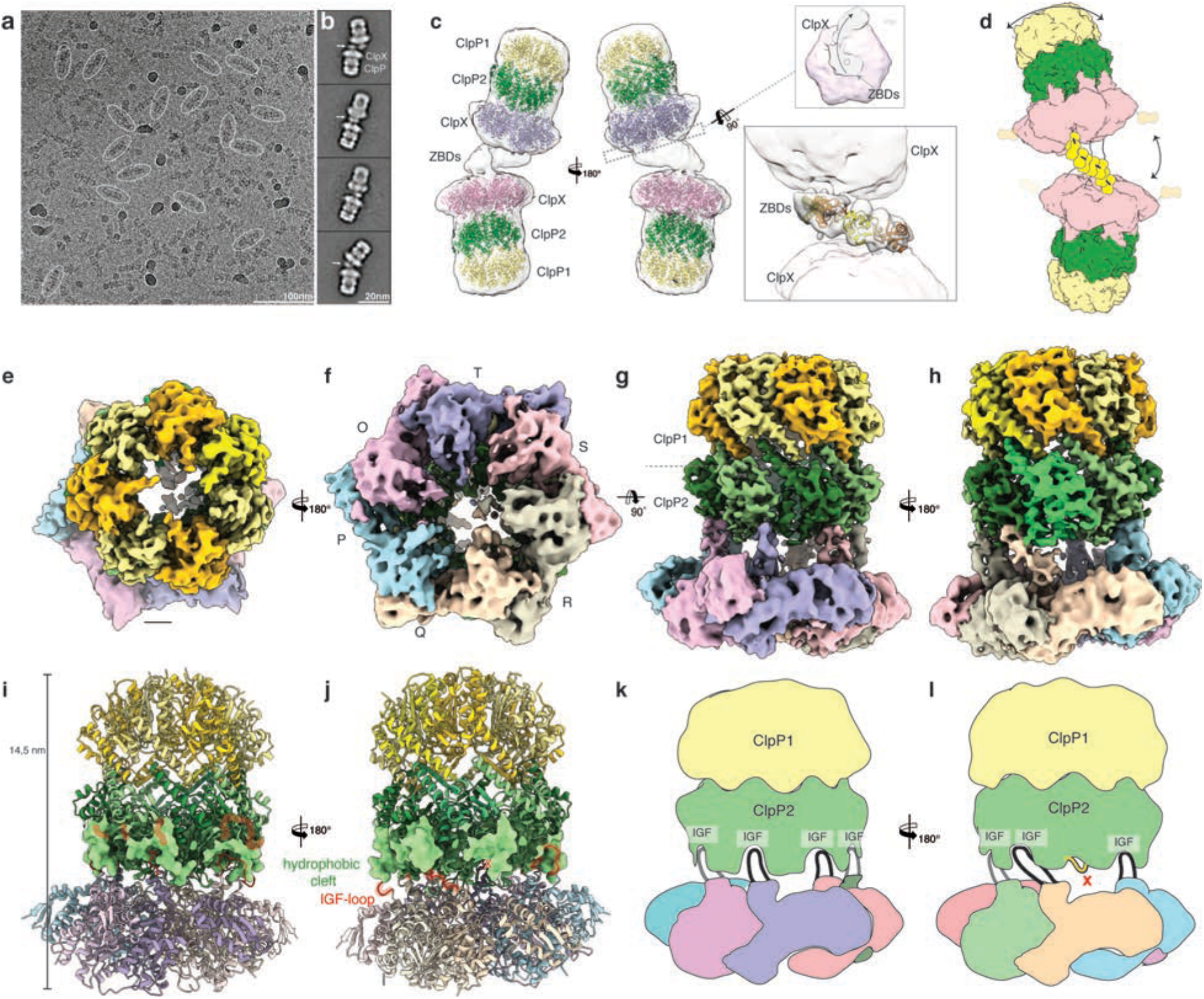
Cryo-EM structure of the ClpXP1/2 protein degradation machinery. **a)** Typical low-dose cryo-EM micrograph of the ClpXP1/2 dimer from *L. monocytogenes*. Some particles are highlighted with ovals. Scale bar, 100nm **b)** Typical reference-free 2D class averages. Arrows indicate additional densities corresponding to ZBDs at the interface between two ClpX hexamers. Scale bar, 20nm **c)** Ribbon Model of ClpP1 (yellow), ClpP2 (green) and ClpX (orange) superimposed with the cryo-EM density map of the ClpXP1/2 dimer (white and transparent). The upper inset shows the complex shown as slice at the position of the axial pore entry of the upper ClpXP1/2 complex. ClpX and ClpX-ZBD densities are colored magenta and gray transparent, respectively. The arrow indicates the spiral arrangement of the ZBD domains. The lower inset shows the manual fitting of four copies of ZBD-dimers (PDB: 1OVX) into the cryo-EM density at the interface between the ClpX hexamers. The dimers are interconnected as proposed in (*24*). **d)** Cartoon depicting ClpXP1/2 dimerization via the ZBD domains of two opposing ClpX hexamers. Arrows indicate the flexibility of the complex. **e-h)** Cryo-EM density of ClpXP1/2 shown from the top (**e**), bottom (**f**) and side (**g,h**). ClpP1 and ClpP2 subunits are colored in khaki, orange and dark, light green, respectively. ClpP2 subunit J is highlighted in mint green. Note that this is the only ClpP2 subunit not interacting with ClpX via an IGF-loop. Each subunit of ClpX is assigned a different color. This color code is maintained throughout the manuscript. **i-j)** Molecular model of ClpXP. The hydrophobic pockets of ClpP2, each spanning two ClpP2 subunits, are shown as surface. The IGF interaction loops are highlighted in red. **k-l)** Cartoon depicting how the ClpX hexamer interacts with the ClpP2 heptamer via the six IGF-loops. Note the extended conformation of IGF-loop of ClpX subunit Q.

The ZBDs are involved in substrate binding and cofactor recognition and were shown to dimerize when expressed as single domain (*23*, *24*). Based on these results it has been previously proposed that the ZBDs of neighboring subunits within a single ClpX hexamer dimerize resulting in a trimer-of-dimer model (*24*). In this model the ZBD dimers interact with the adjacent dimers, creating a ring structure that is aligned with the central channel of ClpX. The structure of the ClpXP1/2 dimer, however, reveals that the ZBDs do not form rings, but arrange in a flexible half-cone spiral with the first and last ZBD dimer positioned directly above or at the rim of the axial pore entry of the upper and lower ClpX hexamer, respectively (Figure 1c, Supplementary Figure 1e). The ZBDs are apparently interacting with the ZBDs from oppositely positioned subunits leading to the cross-linking of the two opposing ClpX hexamers (Figure 1c, d). In total, four ZBD dimers fit into the cryo-EM density (Figure 1c). Because of the limited resolution in this region, however, we cannot determine if the cross-bridges are mediated by single ZBDs that dimerize with ZBDs of the other ClpX or by ZBD dimers that interact with dimers of the other ClpX. The half-cone spiral arrangement together with the high flexibility of the interaction between the two opposing ClpX hexamers (Supplementary Video 1) might be involved in substrate binding and probably even help guiding it into the ClpX pores. Based on these results and the fact that the ZBDs are flexible and not resolved in the crystal structure of ClpX(*25*), we propose that ZBD dimers form stable structures only at the interface between two oppositely positioned ClpX hexamers (Figure 1d).

To obtain a cryo-EM structure at higher resolution, we focused the structural analysis on one ClpXP1/2 subunit in the dimer and solved its structure using the same dataset (Figure 1e-l, Supplementary Fig. 2). The final cryo-EM reconstruction has an average resolution of 3.6 – 4 Å for ClpP1/2 and 6 – 7 Å for ClpX (Supplementary Fig. 2e-g). The overall lower resolution of ClpX indicates that the chaperone is intrinsically more flexible and heterogeneous than the ClpP barrel in the ClpXP1/2 complex. To build a complete atomic model of ClpXP1/2, we fitted a homology model of ClpX and the available crystal structure of ClpP1/2 (PDB-ID 4RYF) into the cryo-EM density and refined the model using Molecular Dynamics Flexible Fitting (MDFF) (*26*).

The structure of ClpXP1/2 reveals that ClpP1 forms the upper homoheptamer of the ClpP barrel, whereas ClpP2 sits below and interacts with ClpX (Figure 1g-l). This arrangement is consistent with the results of previous binding studies showing that ClpX and ClpP1/2 interact exclusively via the ClpP2 ring surface in *Listeria monocytogenes* and *Mycobacterium tuberculosis* ClpP proteases (*27*, *28*).

Interestingly, the ClpX hexamer is not centrally aligned, but slightly tilted by ~11° towards ClpP2. The structure of ClpP1/2 is almost identical to the available crystal structure of apo-ClpP1/2 (PDB-ID 4RYF), indicating that the binding of ClpX does not induce large conformational changes in ClpP1/2 (Figure 5b). In contrast, the interaction with ClpP1/2 has an effect on the overall conformation of ClpX. Whereas the AAA+ domains arrange as a “dimer-of-trimers” in the crystal structure of *E. coli* ClpX (*29*), the structure of ClpXP1/2 shows that the ring of ClpX exhibits pseudo-six-fold symmetry. In addition, the neighboring AAA+ domains are closely packed to each other. This is unlike recent structural studies on substrate-bound AAA+ chaperones that showed a “spiral-staircase” arrangement with one “seam” subunit, which is slightly displaced from the pore (*30*–*32*). The resolution at the nucleotide pocket is not high enough to visualize nucleotides, but the structure reveals that all six ClpX protomers are in the “loadable” conformation (Supplementary Fig. 3). This is in contrast to ClpX with the E183Q mutation in its apo-state (*29*). There, two subunits are in the “loadable” and four are in the “unloadable” conformation. A dynamic interconversion between loadable and unloadable conformations is required to couple ATP hydrolysis by ClpX to mechanical work. However, the arrangement is not a direct consequence of the bound nucleotide or the presence of specific mutations (*25*). To further examine the interaction between ClpP1/2 and ClpX we used hydrogen-deuterium exchange with mass spectrometry (HDX-MS) to monitor the accessibility of residues at the interface. In line with our structural observations, complex formation between ClpP1/2 and ClpX only changes the accessibility of residues of ClpX and ClpP2, but not of ClpP1 (Supplementary Fig. 4). This not only corroborates that ClpX solely interacts with the ClpP2 isoform, but also indicates that ClpX binding does not induce major allosteric conformational changes in the ClpP1 heptamer.

### Symmetry mismatch of IGF-loop interaction

The most interesting part of the structure is the interface between ClpP2 and ClpX, which involves a C6/C7 symmetry mismatch. As predicted by biochemical studies (*8*, *11*, *33*), it is mediated mainly by the flexible IGF loops of ClpX interacting with hydrophobic grooves in ClpP2 (Figure 1g-h). The tilted arrangement of ClpX results in part of the loops interacting stronger with ClpP2 than others (Figure 2a).

**Figure 2.**
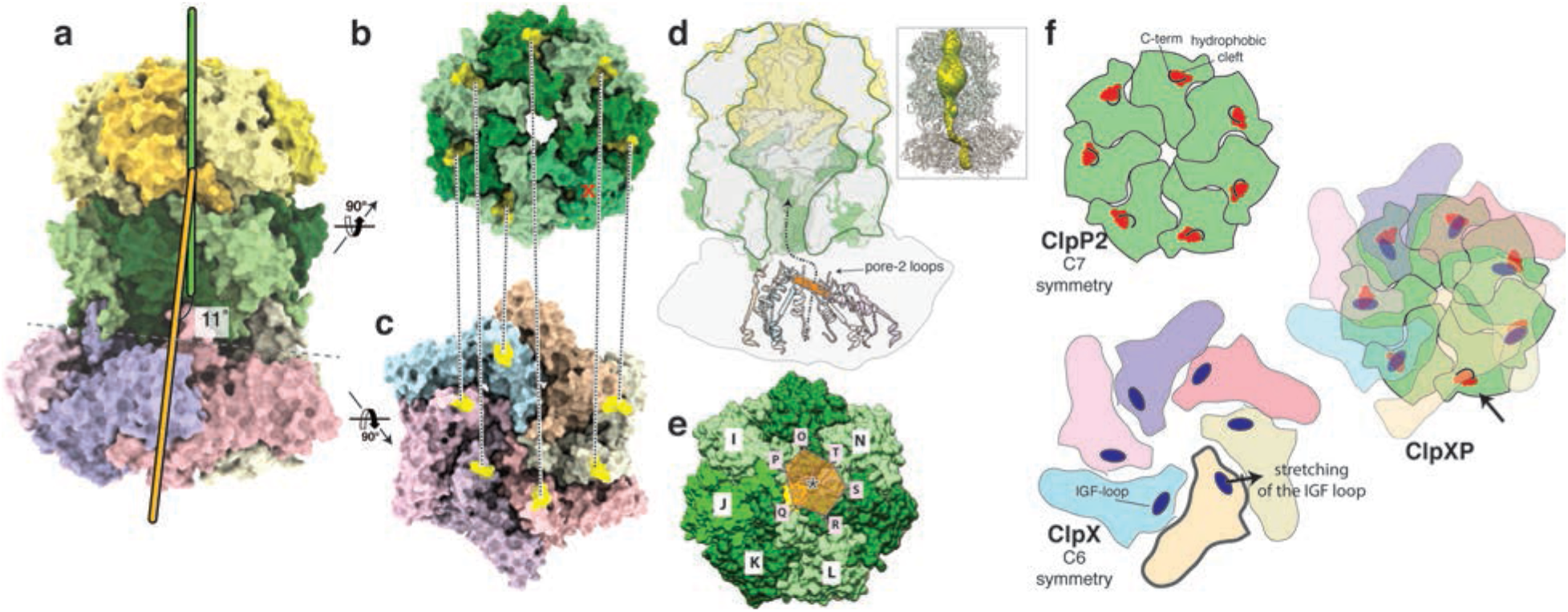
Symmetry mismatch between ClpP1/2 and ClpX. **a)** Molecular model of ClpXP1/2 shown as surface. The symmetry axes of the ClpP tetradecamer and the ClpX hexamer are shown in green and orange, respectively. **b-c)** The ClpP2 heptamer **(b)** and the ClpX hexamer **(c)** are shown from the bottom and the top, respectively, perpendicular to the plane of the ClpP2-ClpX interface. The positions of the IGF-loops and the hydrophobic grooves are highlighted in yellow and connected by dashed lines. **d)** Cut-away view of the ClpP density to visualize its axial pores and central lumen. Secondary structure elements directly prior (residues 170-189) and after the pore-2-loops (residues 202-220) of ClpX are shown in ribbon representation. The pore-2-loops are not resolved in the cryo-EM density and not shown here. In order to indicate the arrangement and positioning of the pore-2-loops, as well as the position of the upper opening of the ClpX channel relative to the ClpP2 pore, a plane was calculated using the Cα atoms of Gly202 as anchor points and depicted here in orange. Note that the plane is tilted and shifted relative to the ClpP channel axis, suggesting a spiral staircase-like arrangement of the pore-2-loops. The dashed line with the arrowhead indicates the pathway of substrate translocation from ClpX towards the ClpP proteolytic chamber. The inset shows the skin surface of the ClpXP pore. For calculation and visualization of the pore, the pore-2- and RKH-loops were modeled using Rosetta (Methods). **e)** Molecular surface of ClpP2 shown from the bottom. Rosetta models of the pore-2-loops of ClpX are shown as ribbons. The black star at the center of the pore-2 plane indicates the positioning of the ClpX channel opening relative to the ClpP channel opening (yellow star). **f)** Schematic model of the ClpX-ClpP2-binding mechanism. Left images depict axial views of the ClpP2 heptamer (green) and the ClpX hexamer (coloring similar to Figure 1) prior assembly of the ClpXP protease. The main interaction elements, the ClpX IGF-loops and ClpP2 hydrophobic grooves are highlighted. The C-terminus of each ClpP2 subunit (black) blocks the respective ClpP2 hydrophobic groove. During binding (right image), ClpX is tilted so that five IGF loops come in close proximity to five hydrophobic grooves without major conformational changes. The sixth IGF loop stretches anti-clockwise to reach the next “free” hydrophobic groove, enabling the symmetry mismatch. Thereby, the six IGF-loops push the C-termini of ClpP2 away and bind tightly to the hydrophobic groove. The remaining “free” ClpP2 hydrophobic groove stays shielded by the respective C-terminus (arrow).

The large domains of the respective ClpX subunit from which the loops protrude are positioned directly below the deep hydrophobic grooves of ClpP2 which are formed at the interface of two subunits. This arrangement allows a direct interaction of the IGF-loops with the opposing grooves. The hydrophobic grooves of ClpP are arranged in a circular manner with seven-fold symmetry and the positions of the ClpX IGF-loops in the complex, perfectly match this arrangement. Interestingly, both rings display similar diameters (Figure 2b-c), except that the IGF-ring remains open at the position of the seventh, free hydrophobic cleft.

Five of the six IGF loops (subunits O, P, R, S, T) display an overall similar arrangement. Due to the symmetry mismatch the large domain of the sixth subunit (subunit Q), is positioned in-between two hydrophobic grooves. The respective IGF-loop, however, still interacts with one of the opposing grooves by adopting an “extended” conformation (Figure 1g-i). The other groove stays empty. Although the distance between the IGF-loop and the “left” or “right” ClpP hydrophobic groove are similar, we only obtained a high-resolution structure with the IGF-loop binding exclusively to the left binding pocket.

To support our structural findings, we performed HDX-MS measurements and mutational studies. Upon complex formation, deuterium uptake of the IGF-loop is strongly reduced (Figure 3, Supplementary Fig. 4) and mutations in the IGF loops of ClpX and the hydrophobic grooves of ClpP2 result in impaired complex formation (Supplementary Fig. 5). This is in line with our ClpXP1/2 structure that demonstrates that the interaction between the IGF loops with the hydrophobic grooves is crucial for complex formation and function.

**Figure 3.**
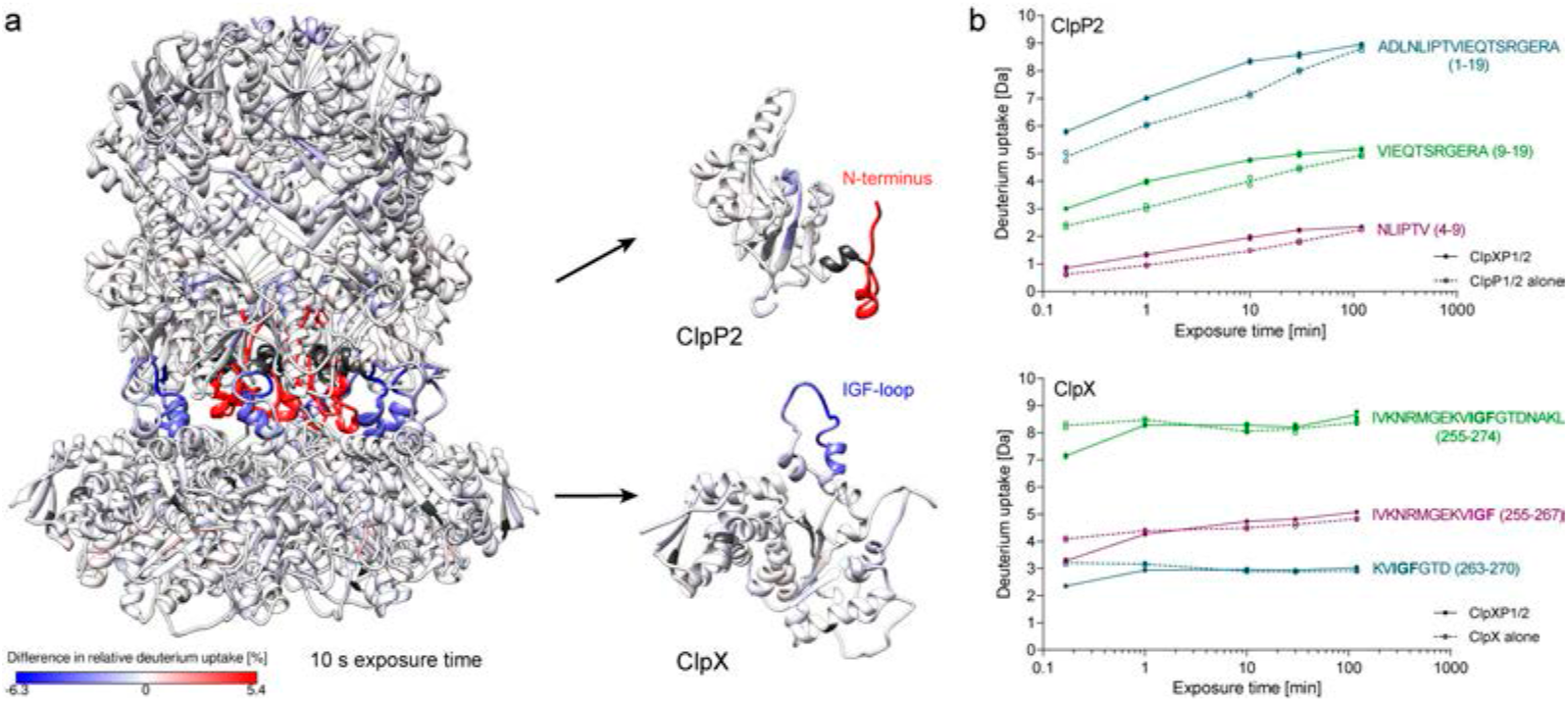
HDX-MS analysis of ClpXP1/2 complex formation. **a)** Difference in relative deuterium uptake after 10 s exposure is mapped on the structure of ClpXP1/2 (left), ClpP2 monomer (right top) and ClpX monomer (right bottom). Increased deuterium uptake upon complex formation is shown in red, decreased deuterium uptake is depicted in blue. Dark gray represents no coverage. **b)** HDX kinetics of exemplary peptides in the N-terminus of ClpP2 (top) and in the IGF-loop of ClpX (bottom). Solid lines and filled circles represent the ClpXP1/2 complex, dashed lines and empty circles represent ClpP1/2 or ClpX. Two independent replicates are shown, lines denote the mean.

Taken together, tilting of the ClpX ring and stretching of one of the IGF-loops is sufficient for the hexameric ClpX to adapt to the seven-fold symmetry of the heptameric ClpP, leaving out one of the binding pockets (Figure 1k-l). Due to multivalence, this results in strong, but at the same time flexible binding, which is likely necessary to accommodate the different conformations of ClpX protomers during ATP hydrolysis and substrate processing (*8*, *19*, *29*).

### The C-terminus of ClpP2 shields the hydrophobic groove prior to ClpX binding

The C-termini of the ClpP2 show two conformations in our structure: a compact conformation that blocks the hydrophobic groove when it does not accommodate an IGF loop, and an extended conformation enlarging the groove when occupied by an IGF loop (Figure 4a). Since the residues of the C-terminus are not conserved (Supplementary Figure 6) and the conformational change is not transmitted to the rest of the protein, an allosteric regulation is rather unlikely. The C-termini probably shield the hydrophobic grooves, when ClpX is not bound and thereby prevent the interaction with other hydrophobic molecules and increase the stability of the protein in a hydrophilic environment.

**Figure 4.**
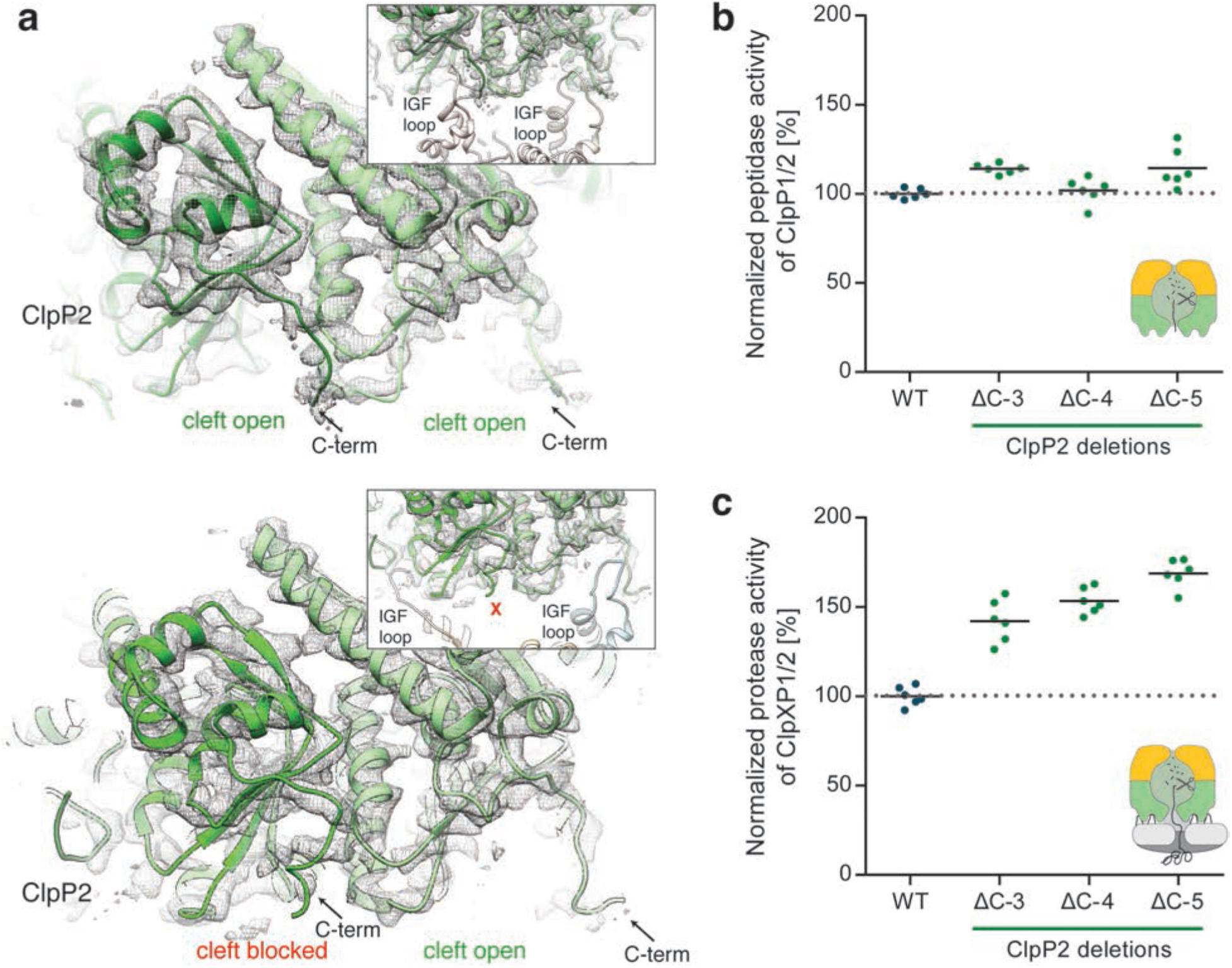
Role of the ClpP2 C-terminus in ClpXP1/2 binding. **a)** Molecular model and cryo-EM density of IGF-loop bound (upper image) and not bound to hydrophobic pockets of ClpP2 (lower image). The insets show the respective IGF-loops in ribbon representation. Arrows indicate the C-terminus of ClpP2. **b)** Peptidase activity of ClpP1/2 with C-terminally truncated ClpP2 (714 nM (ClpP1/2)_14_, 100 μM Ac-Ala-hArg-2-Aoc-ACC). **c)** Protease activity of ClpXP1/2 with C-terminally truncated ClpP2 (0.2 μM (ClpP1/2)_14_, 0.4 μM ClpX_6_, 0.8 μM GFP-SsrA). Data are normalized to the wild type as 100% (n = 6, black lines denote means).

To probe this, we deleted the last three to six amino acids of ClpP2. ClpP1/2^ΔC-6^ precipitated during purification, suggesting that a certain length of the C-terminus is important to protect the hydrophobic groove and facilitate protein stability. ClpP2 mutants bearing three to five amino acid deletions were however soluble and exhibited a similar peptidolytic activity as the wild type complex (Figure 4b). Interestingly, in protease assays requiring the binding of ClpX, the activity increased with a growing number of amino acid deletions in comparison to the wild type complex (Figure 4c). We interpret this result such that when the C-termini are shorter more complexes are formed because ClpX can easier access the hydrophobic grooves via the IGF-loops. Indeed, in line with this finding the C-termini of most ClpPs which were shown to interact with ClpX are shorter in length (Supplementary Figure 6).

### N-termini of ClpP2 and pore-2 loops of ClpX regulate the entry portal

ClpX is not only tilted, but also laterally shifted respective to ClpP2 (Figure 2a, d, e). Such an arrangement has also been described for other complexes that display a symmetry mismatch (*34*–*36*). In the case of ClpXP1/2, this results in a misalignment of the central channels of ClpP and ClpX, creating in a twisted translocation channel with a constriction site at the interface between ClpP2 and ClpX (Figure 2d). At this position, the N-terminal loops of ClpP2 and pore-2 loops of ClpX interact with each other. These interactions are expected to be even more dynamic than the flexible contacts mediated by the IGF loops, and coupled to ATP-hydrolysis (*8*, *11*, *37*). Indeed, the densities corresponding to the N-terminal loops of ClpP2 and pore-2 loops of ClpX are very weak indicating a higher degree of flexibility in this region of the complex (Supplementary Figures 7,8).

Different conformations of the ClpP N-terminal loops have been previously identified in crystal structures of apo and ADEP-bound ClpPs (*9*, *38*, *39*). In the apo *E. coli* ClpP structure, on the apical side of the ClpP barrel the N-termini are in the “down” conformation, opening one axial pore of the barrel. On the basal side six of the N-termini are in the “up” conformation, with the loops moving out of the axial pore, thereby covering and closing it. It was speculated that the six ClpP N-termini in the “down” conformation would open to match the six-fold symmetry of ClpX and the seventh non-interacting N-terminus would stay in the “down” conformation upon binding to the chaperone. However, in the ADEP-bound structure of *E. coli* ClpP all loops point upwards while they are not resolved in a *B. subtilis* ClpP ADEP structure having made general conclusions difficult so far (*38*, *39*).

In our cryo-EM structure, residues 6 to 17 are not resolved, but the rest of the density reveals that all seven N-termini of ClpP2 (the apical side of the barrel facing the chaperone) adopt the “up”-conformation resolving the controversy about their positioning and the accessibility of the pore (Supplementary Fig. 7). The cryo-EM structure demonstrates that the interaction site between the ClpP2 N-termini and the ClpX pore-2 loops is not shielded and freely solvent accessible. In addition, the N-termini undergo a conformational change upon complex formation and adopt the “up” conformation, by which the protein backbone likely gets more solvent exposed and/or flexible. In line with this, deuteration of the ClpP2 N-terminus increased after complex formation (Figure 3, Supplementary Fig. 4). This observation is also supported by reported synchrotron hydroxyl radical footprinting data showing that ClpA binding enhanced the modification rate of an N-terminal peptide of ClpP, pointing towards a higher solvent accessibility (*40*).

### ClpP activation mechanism by ClpX

Previous crystal structures of ClpP in its apo-form, i.e. without ClpX or compound bound, revealed three different conformational states of the protein: “compressed”, “compact” and “extended” (*41*–*45*). The catalytic triad of the peptidase is only intact in the extended state, suggesting that this is the only active state. ADEPs, that bind to the same site on ClpP as the IGF loops, can induce the transition from the compressed to the extended conformation (*13*). In addition, a ~90° rotation of Tyr63 in the hydrophobic pocket results in the widening of the axial pore by 10 – 15 Å. A mutation of this residue to alanine has the same effect (*46*). This “open” extended conformation of ClpP deregulates the protein. Instead of only processing short peptides of five to six residues, it is now capable to degrade large unfolded polypeptides that otherwise could not be processed in the absence of the chaperone (Figure 6d-e) (*38*, *40*, *47*). It has been speculated that the mechanism of ClpP activation by ClpX would imply similar conformational changes (*15*, *46*). The ClpXP1/2 structure demonstrates that this is not the case. ClpP is in the active extended conformation which is very similar to its conformation in the apo-state (Figure 5a, b). Despite the S98A mutation, the catalytic triad is aligned and in its active conformation (Figure 5d). The ClpP1-P2 heptamers are interconnected via typical interactions of antiparallel β9 strands, characteristic for the “extended” active conformation (*41*) (Figure 5e). Importantly, the axial pore of ClpP is not widened, when compared to the crystal structure of the *B. subtilis* ADEP-bound ClpP (Figure 5c, Supplementary Video 2). A comparison of the interface between the IGF-loop and ADEP with the hydrophobic ClpP pocket reveals that both interact with the same non-polar residues including Ile28, Leu49, Tyr63, Phe83, Ile90, Leu115 (Figure 6a-c). However, binding of ClpX does not induce the rotation of Tyr63 (Figure 6c), which is key to opening the pore. Thus, despite the fact that ADEPs and ClpX share the same binding sites, ClpX does not induce the conformational changes resulting in the opening of ClpP. Instead, binding does not induce any major conformational changes and the diameter of the ClpP channel is sufficient to accommodate the unfolded peptides that are threaded into the ClpP pore by the chaperone to be processed sequentially within the chamber of the peptidase (Figure 6f).

**Figure 5.**
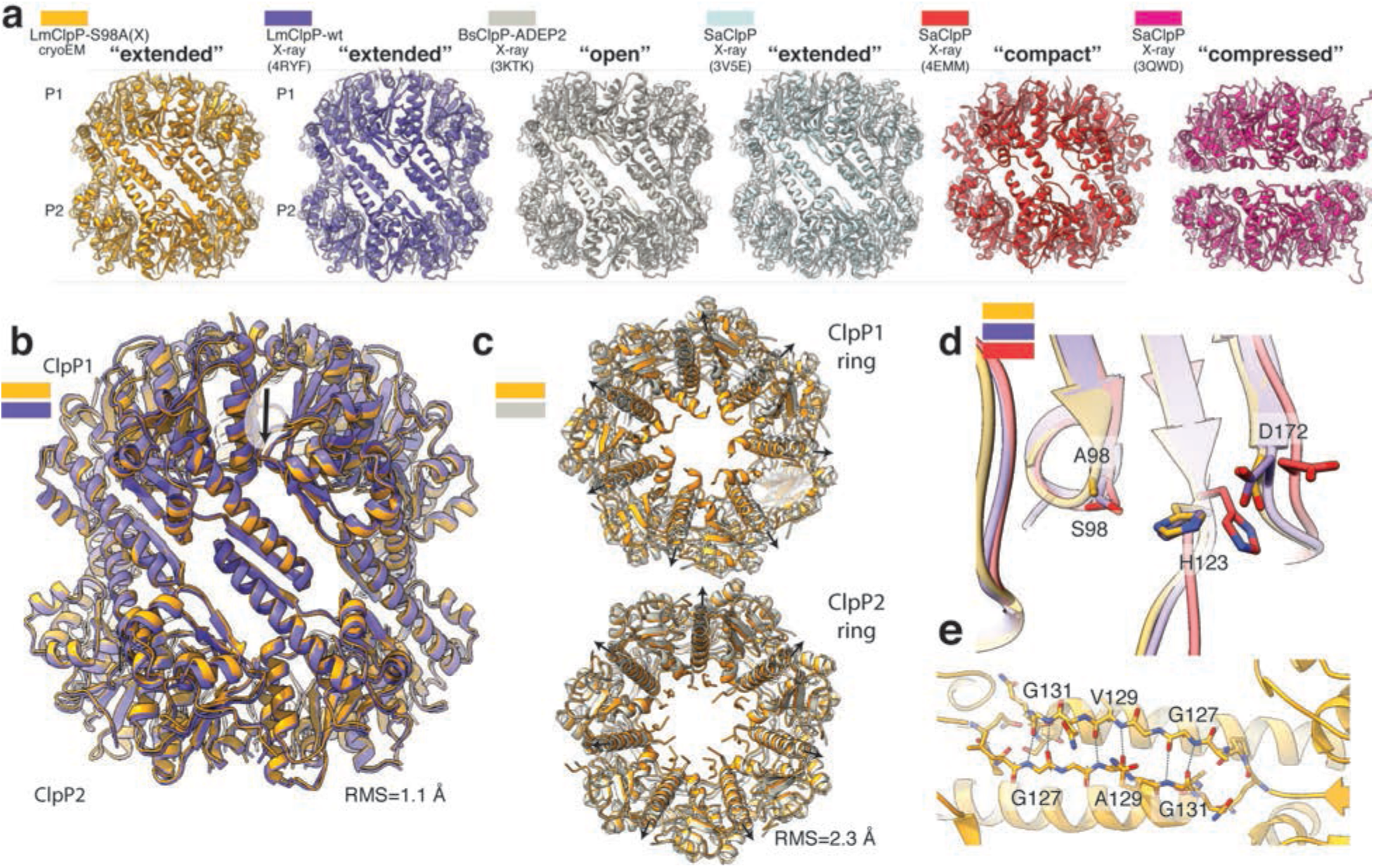
Comparison of ClpX-bound ClpP1/2 with available structures of active and inactive ClpP. **a)** Side view of the structure of ClpX-bound LmClpP1/2 (gold) and the crystal structures of LmClpP1/2 in the extended active state (PDB4RYF) (purple), *Bacillus subtilis* ClpP (BsClpP) in complex with ADEP2 in the extended open active state (PDB 3KTK) (gray), *Staphylococcus aureus* ClpP (SaClpP) in the extended active state (PDB 3V5E) (cyan), SaClpP in the compact inactive state (PDB 4EMM) (red) and SaClpP in the compressed inactive (PDB 3QWD) (purple) conformation are shown in ribbon representation. **b)** Structural superposition of ClpX-bound and unbound (PDB 4RYF) LmClpP1/2. The low R.M.S.D suggests that binding of ClpX to ClpP1/2 does not induce large conformational changes to ClpP1/2. **c)** Structural superposition of ClpX bound ClpP1/2 heterocomplex and ADEP2-bound ClpP homocomplex (PDB 3KTK) shown in top- and bottom view. Black arrows indicate the characteristic opening of the ClpP pore upon ADEP binding. **d)** Superposition of the catalytic residues S98 (S98A), H123 and D172 (N172) in ClpX-bound LmClpP1-S98A/P2-S98A, LmClpP1/2 (extended active state) (PDB 4RYF), SaClpP (compact inactive state) (PDB 4EMM). Note that despite the S98A mutation, the catalytic residues of ClpX-bound LmClpP1/P2 adopt the active conformation. **e)** Opposing subunits of ClpX-bound ClpP1 and ClpP2 rings interact via an antiparallel β-sheet.

**Figure 6.**
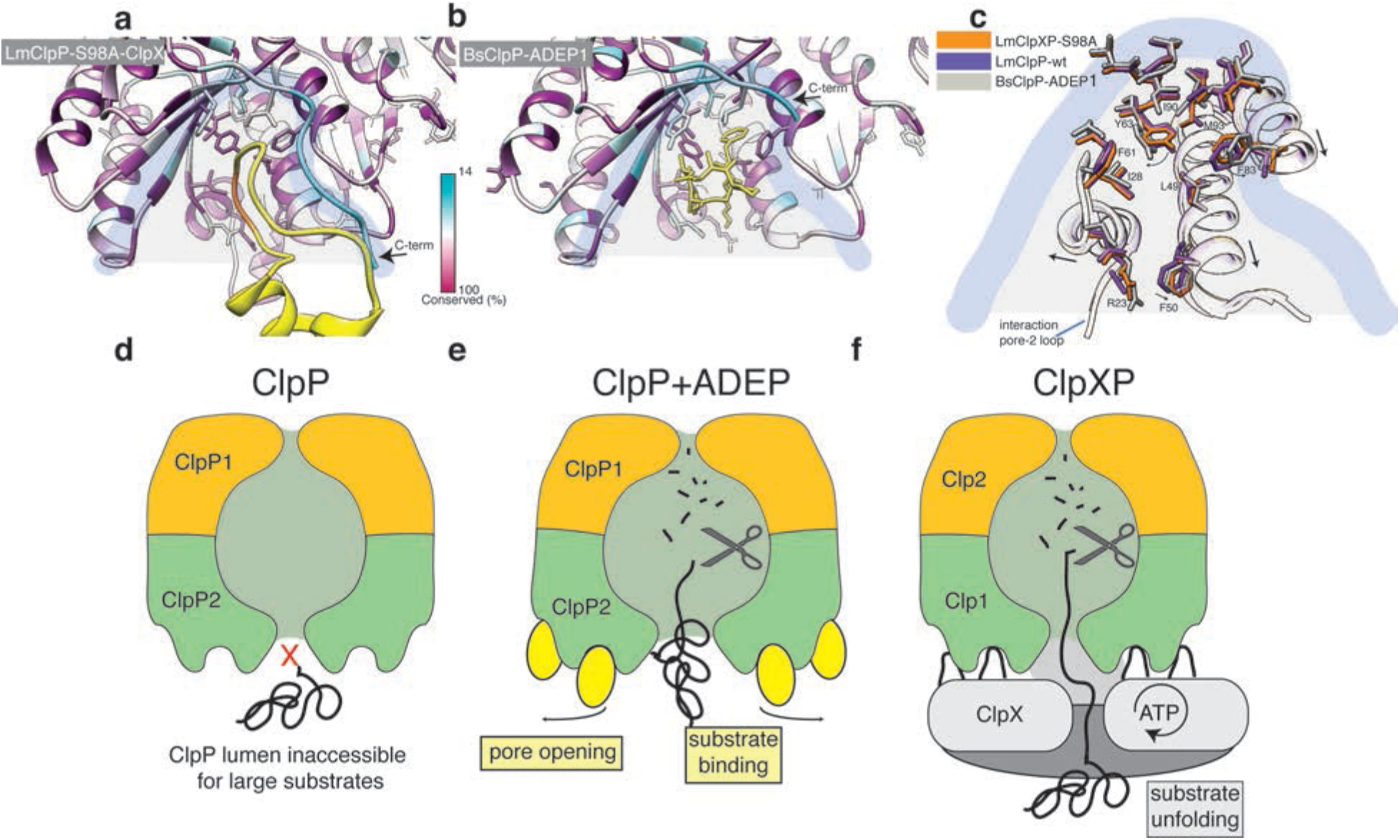
ClpX binds to ClpP in a similar manner like ADEP, but does not induce ClpP pore widening. **a)** Local molecular interactions at one of the seven binding pockets between ClpP and the IGF-loop of ClpX. Residues of ClpP are colored by sequence conservation. The IGF-loop is shown in yellow with the IGF residues highlighted in orange. **b)** Interface of ADEP2 (yellow) with BsClpP (PDB 3KTI) (colored by conservation). **c)** Structural superposition of the binding pockets of ClpX-bound LmClpP2-S98A, ADEP1-bound BsClpP (PDB 3KTI) and “free” LmClpP2 (PDB 4RYF). Arrows indicate changes upon ADEP binding. **d-f)** Regulation of ClpP by ClpX and ADEP. The central pore of the ClpP protease is closed and entry of folded proteins into the proteolytic chamber is not allowed **(d).** ADEP binding to the binding pockets of ClpP induces pore opening. The proteolytic chamber is now accessible for unfolded proteins, leading to unregulated protein degradation and cell death. **(e)** ClpX binds in the same hydrophobic pockets on ClpP but does not induce pore opening. ClpP and ClpX form a continuous pore instead, with ClpX unfolding target proteins and forwarding them to the proteolytic chamber of ClpP for degradation in a regulated manner **(f)**.

## Concluding remarks

The structure of ClpXP1/2 from *L. monocytogenes* allowed us to answer a plethora of long-standing questions regarding complex formation and interaction between the involved subunits. We elucidated the sophisticated interaction between ClpP2 and ClpX that involves a pseudo-six-fold to seven-fold symmetry mismatch at molecular level. We provide the first insights on how ClpX and ClpP together form a continuous twisted translocation channel via interactions between the pore-2-loops of ClpX and the N-termini of ClpP. The C-terminus of ClpP shields the ClpX binding sites prior ClpX binding, which might be important for the future design of ClpP-based antibiotics. The structure surprisingly revealed that although ClpX interacts with the same site on ClpP as the potential antibiotic ADEP, the underlying mechanism of action and structural changes differ considerably. Our data provide intriguing and unexpected insights into the ClpXP architecture, which is essential to understanding the molecular mode of action of this dynamic and highly flexible protein degradation machinery.

## Acknowledgements

We thank O. Hofnagel for assistance in electron microscopy and Dr. M. Lakemeyer for building ClpX homology models. We are grateful to Dr. M. Haslbeck, G. M. Feind and F. Rührnößl for HDX-MS measurements. This work was supported by the Max Planck Society (to S.R.), the European Research Council (FP7/2007-2013) (grant no. 615984) (to S.R.) and the Deutsche Forschungsgemeinschaft (SFB1035) (to S.A.S).

## Author Contributions

S.A.S. and S.R. designed the study. C.G. screened and optimized samples, prepared cryo-EM grids, processed, and analyzed cryo-EM data. D.B. cloned, overexpressed and purified proteins, optimized sample preparation, conducted activity assays and gel filtration measurements, analyzed HDX-MS data. C.G. and F.M. built atomic models. C.G. and D.B. prepared figures, C.G., D.B., S.A.S., S.R. wrote the manuscript. All authors discussed the results.

## Supplementary Figures

**Supplementary Figure 1.**
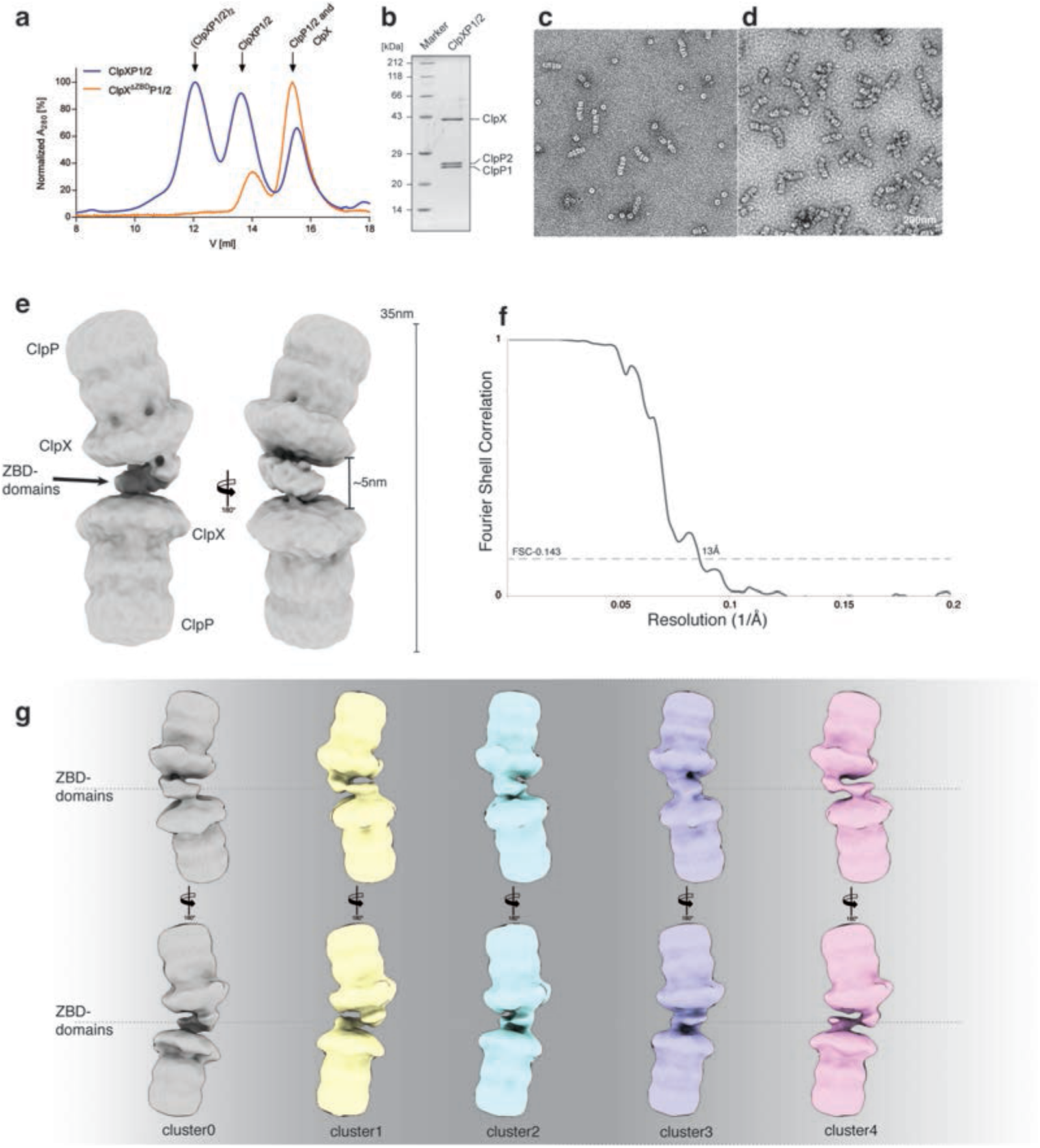
EM analysis of the ClpXP1/2 dimer. **a)** Size exclusion chromatography of ClpX and ClpP1/2 mixtures on a Superose 6 increase 10/300 column. For the EM studies, a sample at 12 mL was taken. Note that the (ClpXP1/2)_2_ peak is absent with ClpX^ΔZBD^. **b)** SDS-PAGE of the isolated (ClpXP1/2)_2_ complex. **c-d)** Subarea of a negative stain EM micrograph of the isolated (ClpXP1/2)_2_ prior **(c)** and after **(d)** crosslinking; Scale bar, 200 nm **e)** Low resolution cryo-EM density of the ClpXP1/2-dimer **f)** Fourier Shell Correlation (FSC) between two independently refined half maps **g)** 3D clustering of the ClpXP1/2-dimer dataset.

**Supplementary Figure 2.**
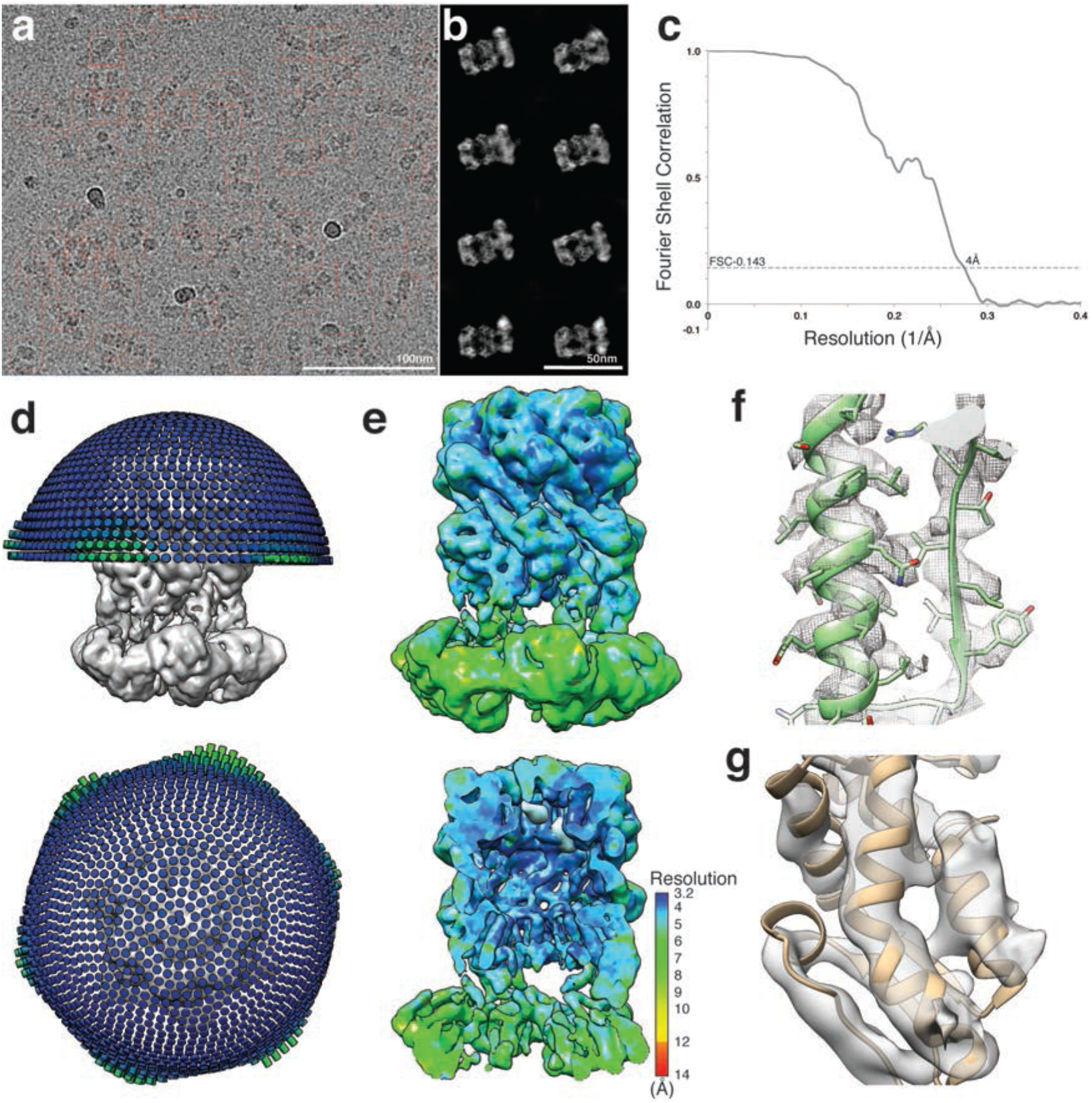
Cryo-EM analysis of ClpXP1/2. **a)** Subarea of a typical low-dose cryo-EM micrograph of ClpXP1/2. ClpXP1/2 particles were selected and extracted from ClpXP1/2-ClpXP1/2 dimers using crYOLO and highlighted in red boxes. Scale bar, 100 nm **b)** Representative reference-free 2D class averages of ClpXP1/2. Scale bar, 100 nm **c)** Fourier Shell Correlation (FSC) between two independently refined half maps **d)** Orientation distribution of the particles used in the final refinement round **e)** Side and cut-off view of the density map colored according to the local resolution **f-g)** Superposition of segments of the molecular model of ClpP **(f)** and ClpX **(g)** with the cryo-EM density (transparent surface).

**Supplementary Figure 3.**
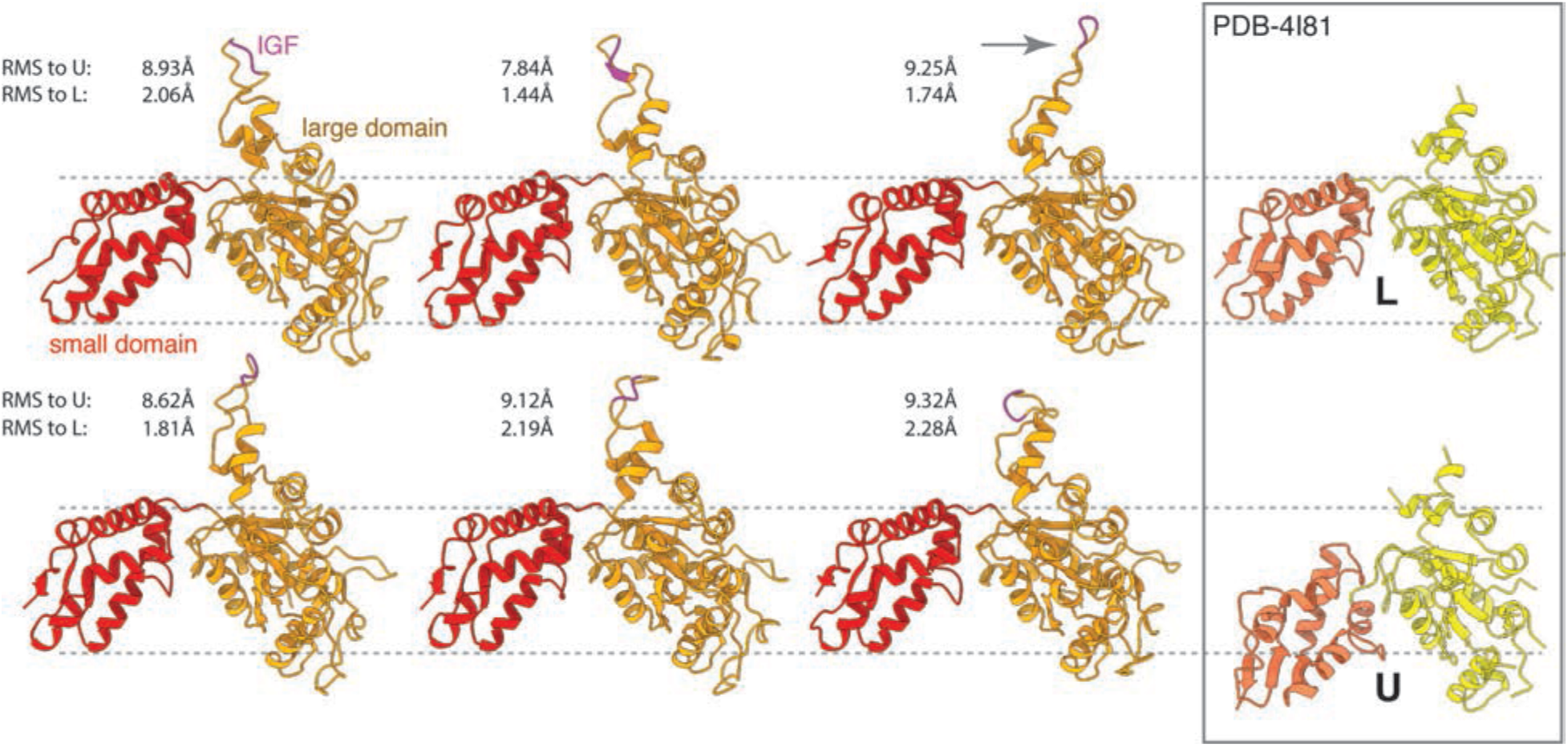
ClpP-bound ClpX subunits adopt a nucleotide-loadable conformation. Molecular models of the six ClpX subunits are shown as ribbon diagrams, with the large domain highlighted in orange and the small domain in red. Note the high similarity between the ClpX subunits, except the arrangement of their IGF-loops. The gray arrow indicates the IGF-loop of subunit Q that adopts an “extended” conformation. The inset shows nucleotide-loadable (L; upper image) and unloadable (U; lower image) subunits of ATP_Υ_S-bound EcClpX (PDB 4I81). Structural comparison of the six ClpX subunits with the L and U subunit of EcClpX (note the respective RMSD values of the Cα atoms) indicate that all subunits of ClpP-bound ClpX adopt a loadable conformation.

**Supplementary Figure 4.**
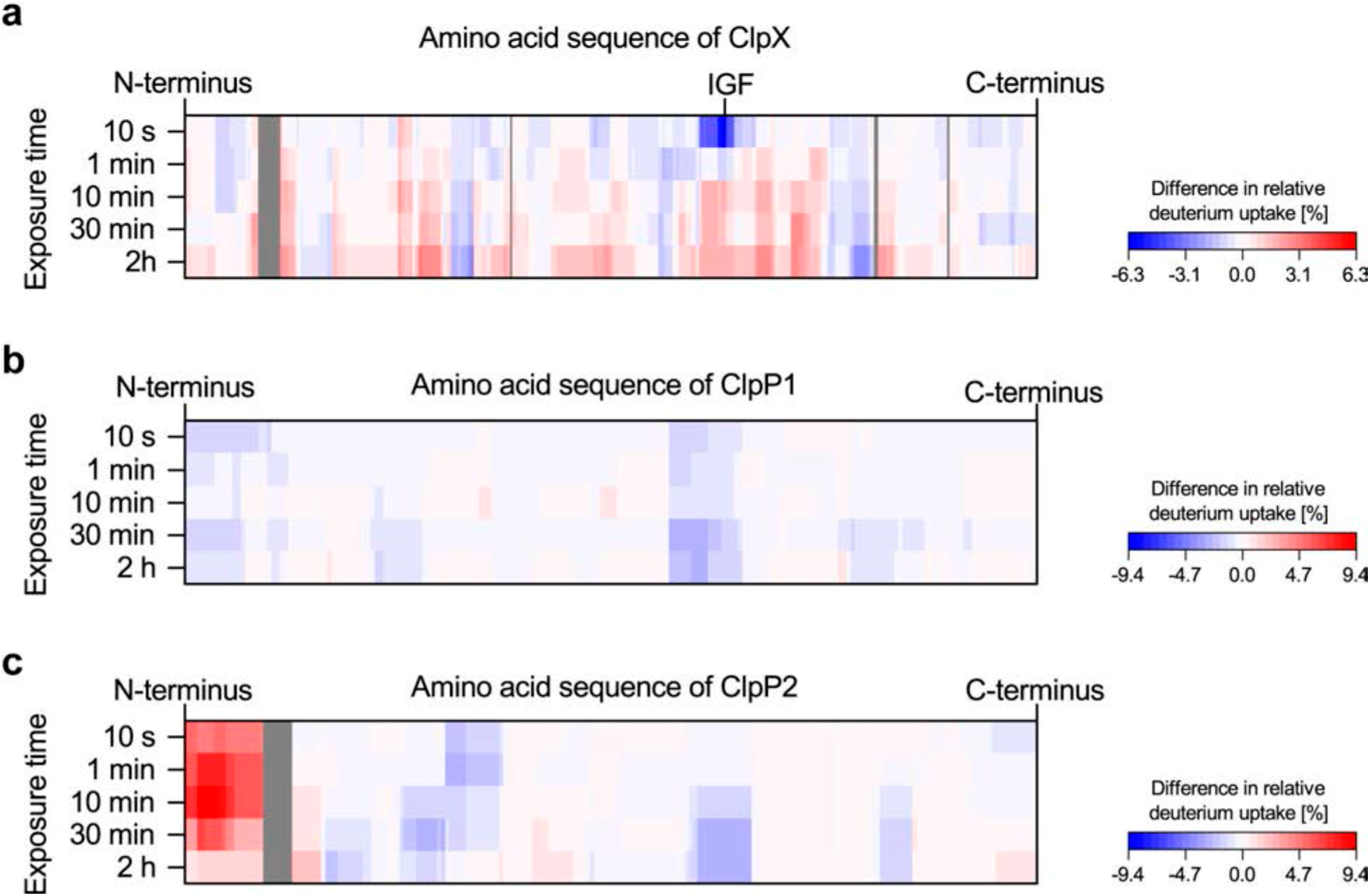
HDX-MS of the ClpXP1/2 complex. Changes in deuterium uptake after complex formation are mapped on the amino acid sequence of **a)** ClpX, **b)** ClpP1 and **c)** ClpP2 for the respective exposure times. Increased deuterium uptake upon complex formation is shown in red, decreased deuterium uptake is depicted in blue. Dark gray represents no coverage. Please refer to the Methods section for the calculation of the relative deuterium uptake values. Averages of two independent measurements are shown.

**Supplementary Figure 5.**
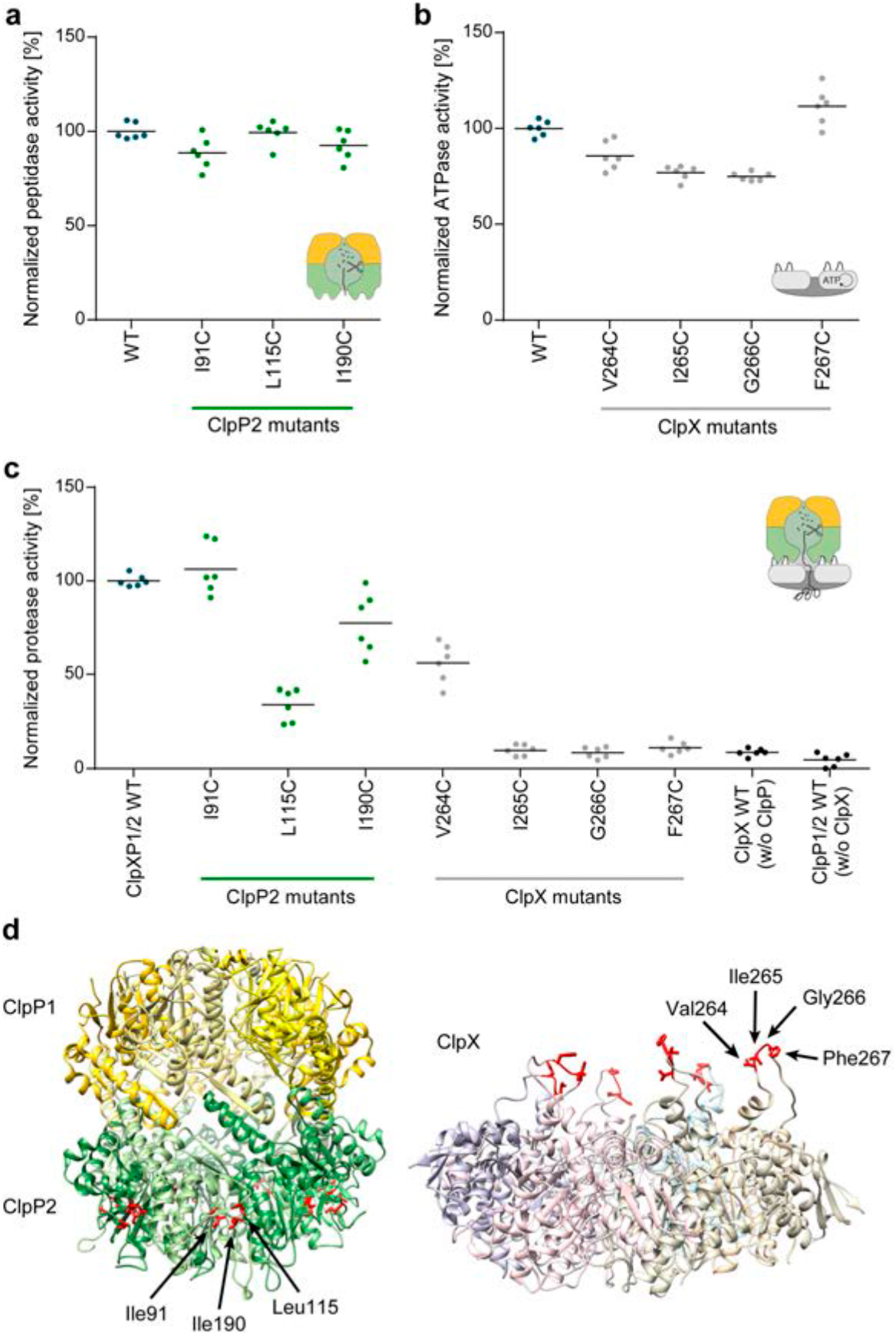
Activity assays of ClpX and ClpP2 mutants of the IGF-loop/hydrophobic groove interface. **a)** Peptidase activity of ClpP1/2 with respective ClpP2 mutants (0.71 μM (ClpP1/2)_14_, 100 μM Ac-Ala-hArg-2-Aoc-ACC). **b)** ATPase activity of ClpX mutants (0.33 μM ClpX_6_, 20 mM ATP). **c)** Protease activity of ClpXP1/2 with ClpP2 and ClpX mutants (0.2 μM (ClpP1/2)_14_, 0.4 μM ClpX_6_, 0.8 μM GFP-SsrA). Data are normalized to the wild type as 100% (n = 6, black lines denote means). **d)** Mapping of the ClpP2 and ClpX mutations on the protein structure. Mutation sites are shown with red sticks.

**Supplementary Figure 6.**
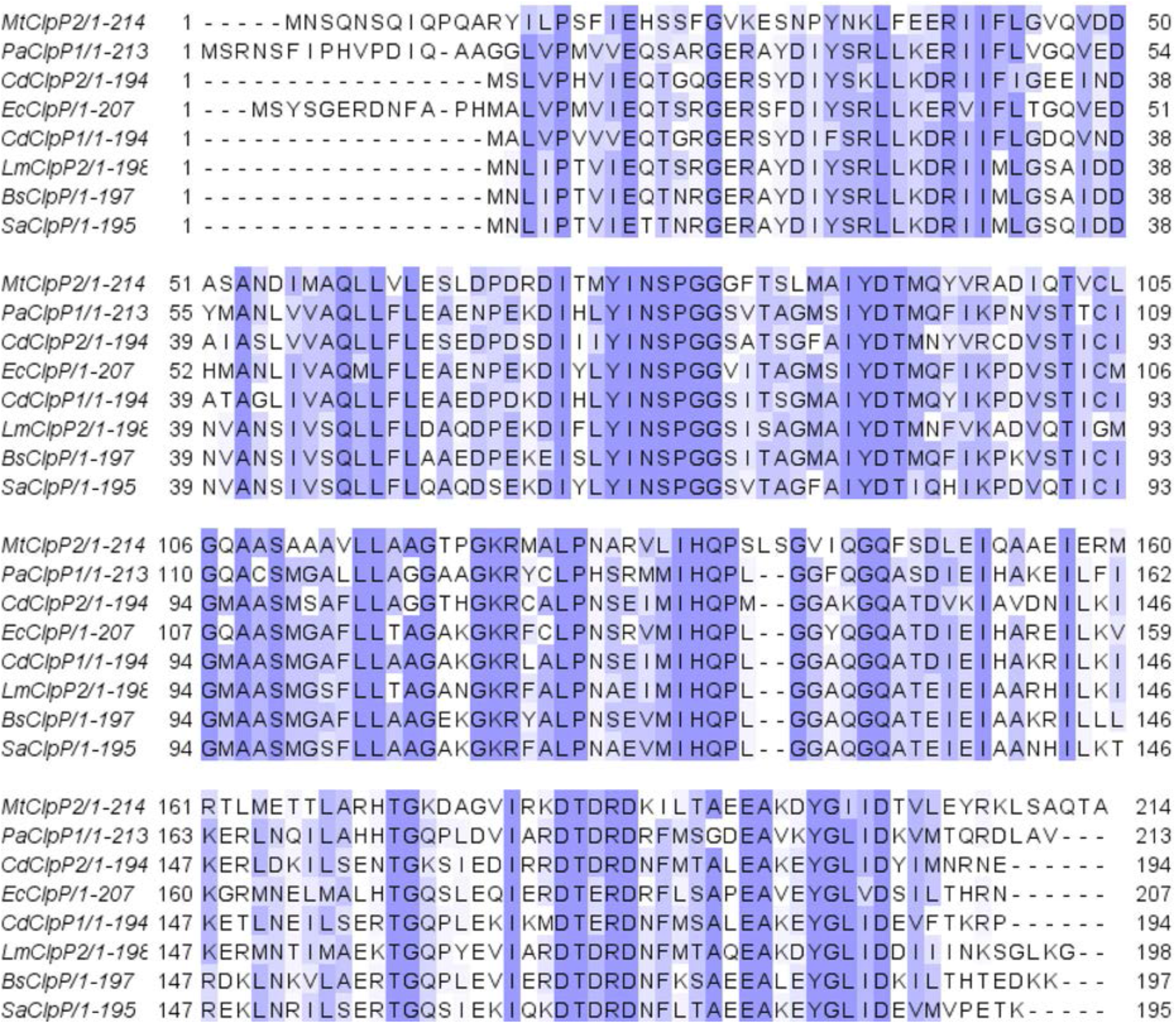
Alignment of ClpP sequences. Mt = *Mycobacterium tuberculosis* (ClpP2), Pa = *Pseudomonas aeruginosa* (ClpP1), Cd = *Clostridium difficile* (ClpP1 und ClpP2), Ec = *Escherichia coli*, Lm = *Listeria monocytogenes* (ClpP2), Bs = *Bacillus subtilis*, Sa = *Staphylococcus aureus*

**Supplementary Figure 7.**
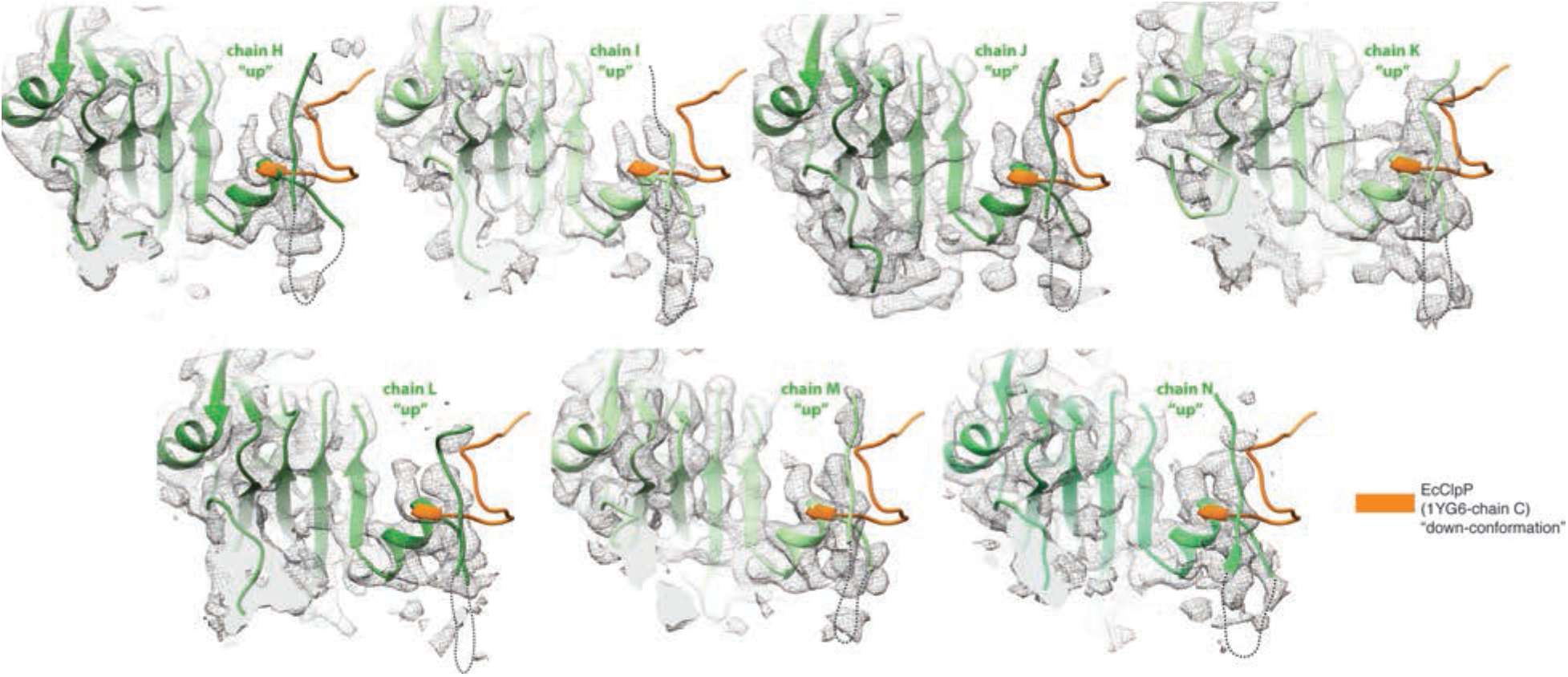
The N-terminal loops of ClpX-bound ClpP2 subunits adopt the “up” conformation. Cryo-EM density map (mesh) with the molecular model highlighting the N-terminal domain of the seven ClpP subunits, with the N-terminal loop modeled in the “up” conformation. Dashed lines indicate residues not resolved in the cryo-EM density. The molecular model of a subunit of EcClpP (PDB 1YG6) with the N-terminus in the “down” conformation (orange) is also shown superimposed.

**Supplementary Figure 8.**
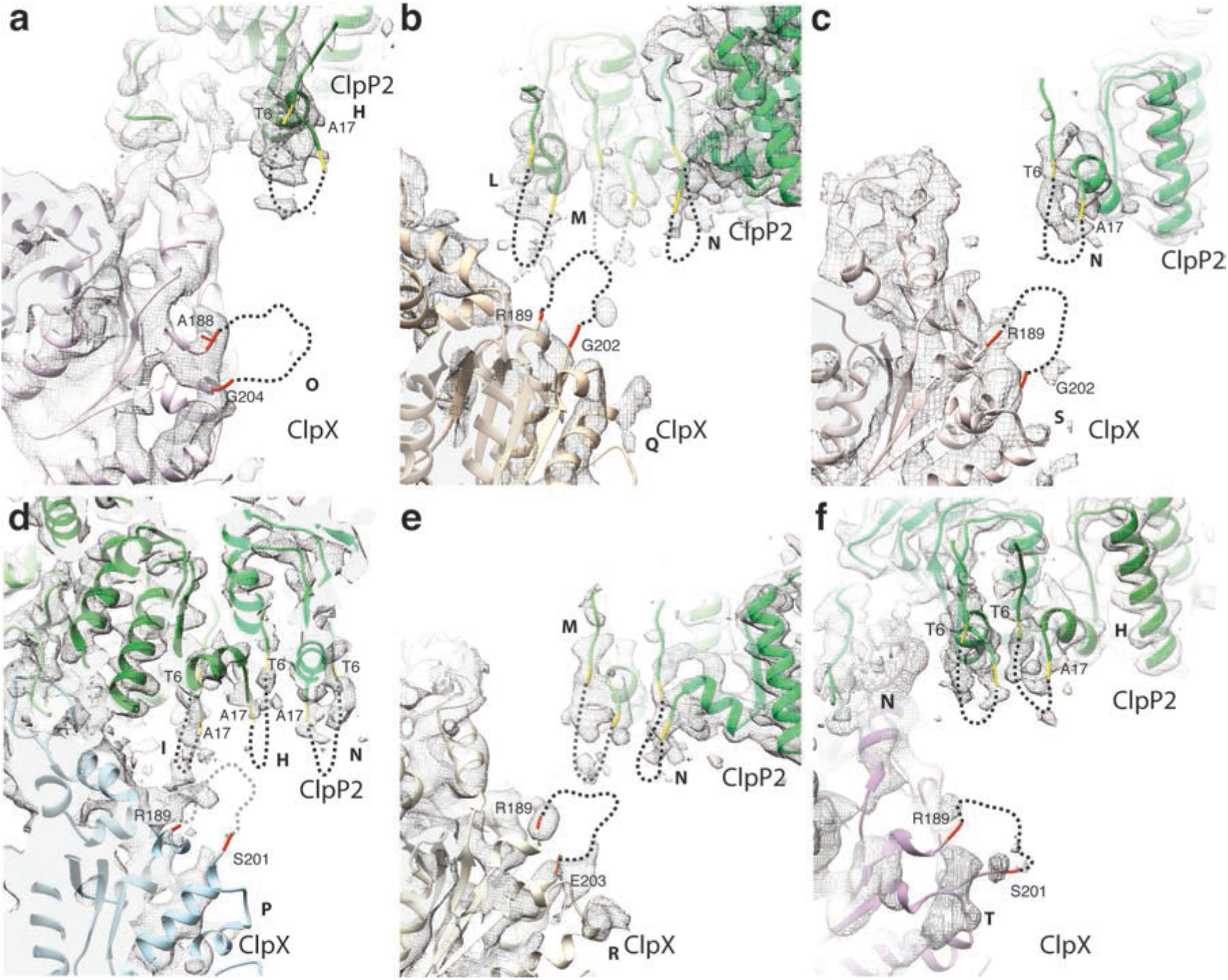
Possible interactions between pore-2-loops of ClpX with the N-termini of ClpP. **a-f)** Cryo-EM density map with the molecular model, highlighting the interaction area between the pore-2-loops of ClpX and the N-termini of ClpP. The six pore-2-loops of ClpX and residues 7-16 of the N-termini of seven subunits of ClpP2 are not resolved. Possible arrangements of these regions are indicated by dashed lines, based on their anchor points and number of residues. Note that pore-2-loops of chains Q and P point into a cleft formed by three ClpP N-termini **(b,c)**. This topological analysis also suggests that pore-2-loops of chains O and T **(a,f)** do not show any interactions with N-termini of ClpP. However, an unusual stretched conformation of these pore-2 loops towards ClpP cannot be excluded. Pore-2-loop of chain S is positioned in direct proximity with N-terminus of chain N **(c)** whereas pore-2-loop of chain R is positioned between two ClpP N-termini (M and N)**(e)**.

**Supplementary Table 1:**
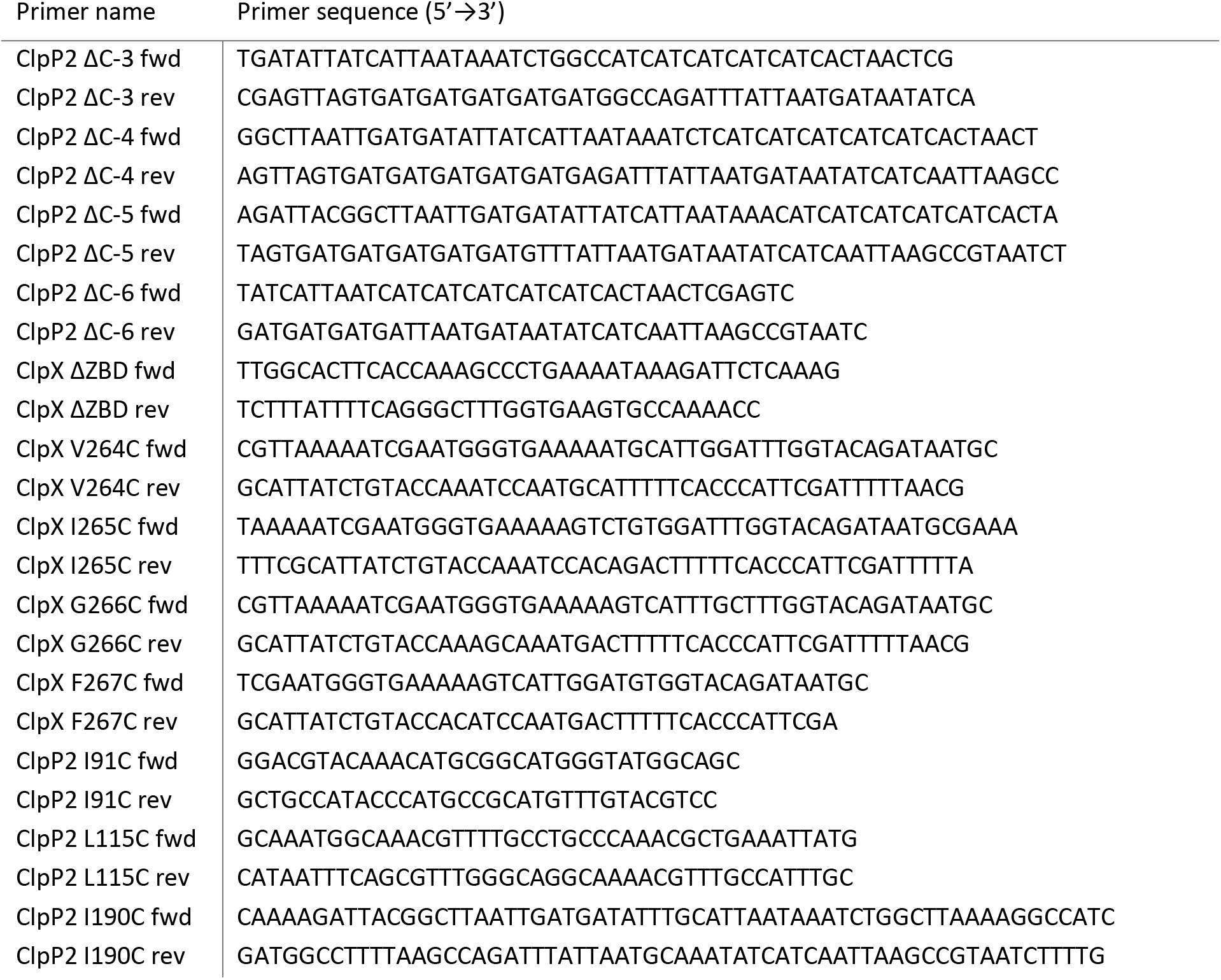
Cloning primers.

**Supplementary Table 2.** Cryo-EM data collection, refinement and validation statistics.

|  | ClpXP1/2 dimer<br>(EMDB-xxxx) | ClpXP1/2<br>(EMDB-xxxx)<br>(PDB xxxx) |
| --- | --- | --- |
| **Data collection and processing** |  |  |
| Magnification | x112,807 | x112,807 |
| Voltage (kV) | 300 | 300 |
| Electron exposure (e-/Å<sup>2</sup>) | 114 | 114 |
| Defocus range (μm) | -0.5 - -3.0 | -0.5 - -3.0 |
| Pixel size (Å) | 1.1 | 1.1 |
| Symmetry imposed | C1 | C1 |
| Initial particle images (no.) | 273,300 | 613,322 |
| Final particle images (no.) | 143,901 | 383,927 |
| Map resolution (Å) | 13 | 4 |
| FSC threshold | 0.143 | 0.143 |
| Map resolution range (Å) | - | 3.2-10 |
| **Refinement** |  |  |
| Map sharpening *B* factor (Å<sup>2</sup>) |  | ClpP, avr. -214<br>ClpX, -240 |
| Model composition |  |  |
| Non-hydrogen atoms |  | 34,656 |
| Protein residues |  | 4,461 |
| Ligands |  | - |
| *B* factors (Å<sup>2</sup>) |  | 132.0 |
| Protein |  | - |
| Ligand |  |  |
| R.m.s. deviations |  |  |
| Bond lengths (Å) |  | 0.0091 |
| Bond angles (°) |  | 1.19 |
| Validation |  |  |
| MolProbity score |  | 1.94 |
| Clashscore |  | 10.45 |
| Poor rotamers (%) |  | 0.51 |
| Ramachandran plot |  |  |
| Favored (%) |  | 94.11 |
| Allowed (%) |  | 5.8 |
| Disallowed (%) |  | 0.09 |

**Video 1**

**Flexibility of ClpXP1/2 dimers in 2D.**

**Video 2**

**Structural comparison between ClpX-bound and ADEP-bound ClpP.** The video shows a simple linear interpolation between ClpX-bound ClpP1/2 (cryo-EM) (extended active conformation) and the available crystal structure of *B. subtilis* ADEP2-bound ClpP (PDB 3KTK (*39*)) (extended active open conformation), first along the ClpP1 and finally along the ClpP2 face (see Figure 5c). Note the widening of the pore in the ADEP-bound structure.

## Methods

### Cloning

The cloning of pETDuet-1_ClpP1/2 and pET300_ClpX were described previously (*1*, *2*). ClpX and ClpP1/2 point mutants, ClpP1/2^ΔC-3^, ClpP1/2^ΔC-4^ and ClpP1/2^ΔC-5^ were generated using the QuikChange™ technology. For ClpP1/2^ΔC-4^ and ClpP1/2^ΔC-5^, the pETDuet-1_ClpP1/2^ΔC-3^ plasmid was used as a template. ClpP1/2^ΔC-6^ and ClpX^ΔZBD^(E183Q) were obtained with primers containing non-overlapping sequences (*3*). All primers are listed in Table 1.

**Table 1:**
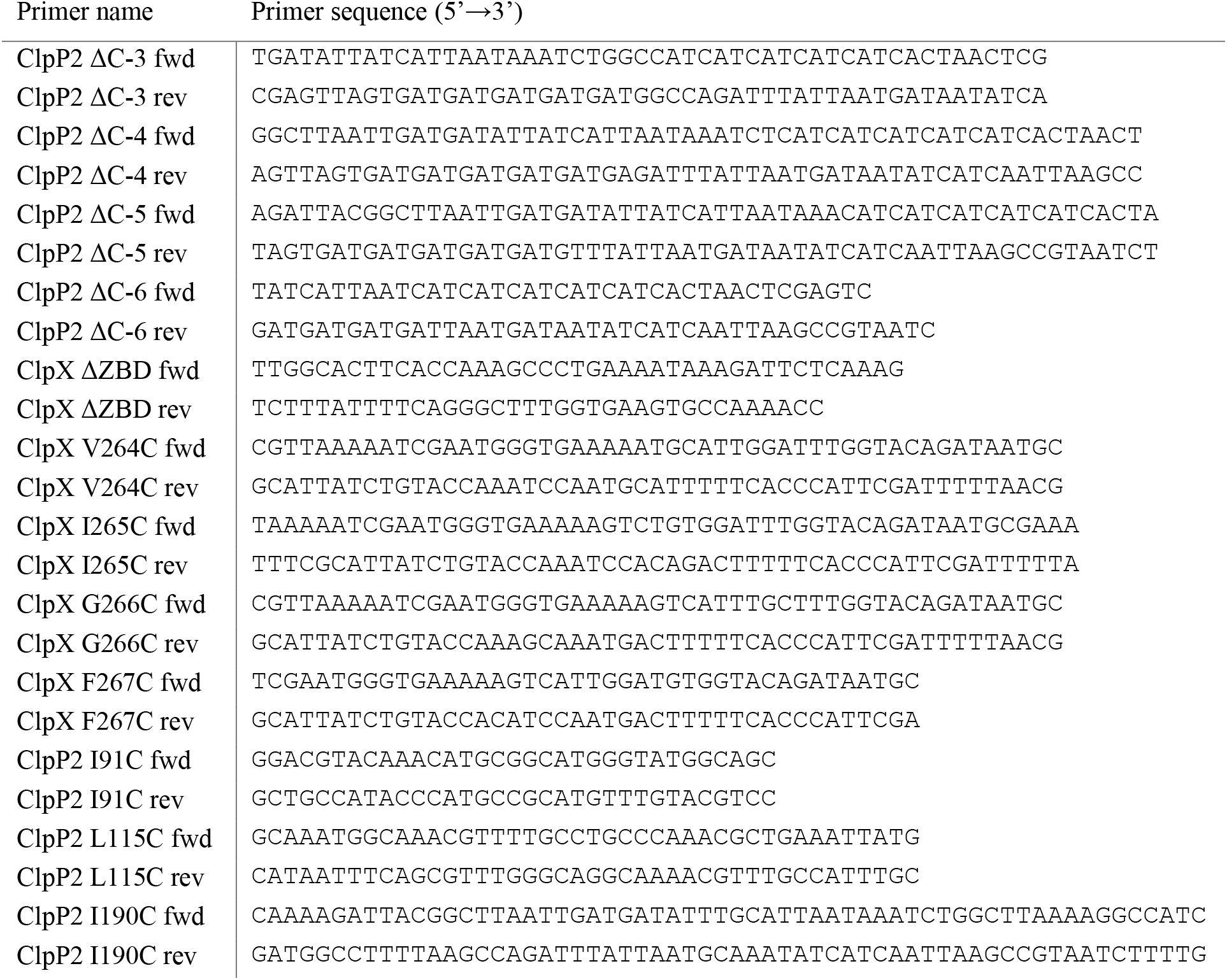
Cloning primers.

### Protein overexpression and purification

ClpP1/2 and its mutants’ variants were overexpressed and purified as follows. The proteins were overexpressed in *E. coli* BL21(DE3) bearing a pETDuet-1 vector with C-terminally Strep-II-tagged ClpP1 and C-terminally His_6_-tagged ClpP2 (*2*). The bacteria were grown in LB medium until OD_600_ 0.6 at 37 °C. Following induction with 1 mM isopropyl-β-D-thiogalactoside (IPTG), the bacteria were incubated at 37 °C for 6 h. After harvest, the cells were sonicated on ice in lysis buffer (20 mM MOPS, 300 mM KCl, 1% CHAPS, 10% glycerol, pH 7.5) and then kept at room temperature during the rest of the purification. The proteins from the cleared cell lysate were captured in a HisTrap HP 5 ml column (GE Healthcare) in His buffers (20 mM MOPS, 300 mM KCl, 10% glycerol, pH 7.5; +40 mM imidazole for washing) using an ÄKTA Purifier 10 system (GE Healthcare). The proteins were eluted by a 15 mL gradient from 40 mM to 300 mM imidazole, and the second elution peak was collected. A subsequent chromatography step was carried out on a StrepTrap HP 5 ml column (GE Healthcare) in Strep buffers (20 mM MOPS, 300 mM KCl, 10% glycerol, pH 7.5; +2.5 mM desthiobiotin for elution). A final gel filtration was performed on a Superdex200 pg 16/60 column (GE Healthcare) in ClpP SEC buffer (20 mM MOPS, 300 mM KCl, 15% glycerol, pH 7.0). In the case of the cystein-containing mutants, 1 mM TCEP was added to all buffers.

ClpX(E183Q) and ClpX^ΔZBD^(E183Q) were overexpressed in *E. coli* BL21(DE3). An expression construct equipped with an N-terminal His_6_-tag and a TEV cleavage site in pET300 vector was used (*2*). The bacteria were grown in LB medium to OD_600_ 0.6 at 37 °C. After induction with 0.5 mM IPTG, the cells were incubated overnight at 25 °C. After harvest, the cells were resuspended in ClpX lysis buffer (25 mM HEPES, 200 mM KCl, 1 mM DTT, 0.5 mM ATP, 5 mM MgCl_2_, 10 mM imidazole, 5% glycerol, pH 7.6) and lysed by ultrasonication. The cleared cell lysate was loaded on a 5 mL HisTrap HP column (GE Healthcare). The column was washed with ClpX wash buffer (25 mM HEPES, 200 mM KCl, 1 mM DTT, 5% glycerol, 40 mM imidazole, pH 7.6). The protein was eluted with ClpX elution buffer (25 mM HEPES, 200 mM KCl, 1 mM DTT, 5% glycerol, 300 mM imidazole, pH 7.6). The protein fractions were pooled, 1 mM EDTA and TEV protease [1.25 mg for ClpX(E183Q) and 3.75 mg for ClpX^ΔZBD^(E183Q)] were added and the reaction mixture was incubated at 10 °C overnight. Complete TEV cleavage was verified by intact-protein mass-spectrometry. The protein solution was loaded on a Superdex200 pg 16/60 column (GE Healthcare) and eluted in ClpX SEC buffer (25 mM HEPES, 200 mM KCl, 1 mM DTT, 0.5 mM ATP, 5 mM MgCl_2_, 5% glycerol, pH 7.6). ClpX(WT), ClpX(V264C), ClpX(I265C), ClpX(G266C) and ClpX(F267C) were overexpressed and purified similarly with the following modifications: the buffers contained 1 mM TCEP instead of DTT, and the ClpX wash buffer and ClpX elution buffer contained additionally 0.5 mM ATP and 5 mM MgCl_2_. The TEV digestion step was omitted.

N-terminally Strep-II-tagged eGFP with a C-terminal SsrA tag (AGKEKQNLAFAA) was overexpressed in *E. coli* SG1146a (Δ*clpP*) using pET55-Dest expression vector and purified by affinity chromatography and gel filtration as described previously (*2*, *4*).

Creatine kinase (product no. 10 127 566 001), lactate dehydrogenase (product no. 10 128 155 001) and pyruvate kinase (product no. 10 127 876 001) were purchased from Roche.

### Isolation of the ClpXP complex

4.4 nmol (ClpP1/2)_14_ and 3.3 nmol ClpX_6_ were incubated for 10 min at 37 °C in PZA buffer (25 mM HEPES, 200 mM KCl, 5 mM MgCl2, 1 mM DTT, 0.5 mM ATP, 15% glycerol, pH 7.6). The samples were loaded onto a Superose 6 increase 10/300 column (GE Healthcare) connected to an ÄKTA Purifier 10 system (GE Healthcare) and eluted at 0.2 mL/min flow rate. Samples were taken at 12 mL retention volume for EM and HDX-MS measurements. For cryoEM, the sample was diluted 1:3 with glycerol-free PZA buffer and 0.1% glutaraldehyde was added. The reaction was quenched after 30 s with 2 eq. Tris-HCl. For SDS-PAGE, 4.4 μg protein was loaded on a gel and stained with Coomassie blue after separation.

### Electron microscopy

Sample quality was examined by negative stain EM. Sample from the respective fraction was further diluted to a concentration of 0.01-0.03 mg ml^−1^ and negative stain EM was performed as described previously (*5*). Images were recorded with a JEOL JEM-1400 equipped with a 4K CMOS detector F416 (TVIPS) at a pixel size of 1.84 Å. For cryoEM, 4 μl of cross-linked ClpXP1/2 dimers at a concentration of 0.045 mg ml^−1^ were applied to a glow-discharged quantifoil 2/1 Cu grid with an additional 2nm thin carbon layer and after an incubation time of 45 sec, rapidly plunge-frozen using a CryoPlunge3 (Cp3, Gatan) at 90% humidity. To improve ice quality and thickness distribution, 0.01% Tween-20 was added shortly prior plunging. The quality of the grids was screened with a JEOL JEM 1400 and a FEI Tecnai Spirit, both equipped with a LaB6 cathode and a 4K CMOS detector F416 (TVIPS). A cryoEM dataset was acquired on a FEI Titan KRIOS at 300 kV equipped with spherical aberration corrector and a Falcon III direct detector (linear mode) at a x112,807 magnification (x59,000 nominal magnification), corresponding to a pixel size of 1.1 Å. Each exposure was recorded with a total dose of ~114 electrons/Å^2^ and a total exposure time of 2 sec (frame rate of 50 msec). A total of 3200 micrographs were collected using the EPU software (FEI).

### Image processing and reconstruction

The frames were aligned, averaged and dose-weighted using unblur and sum_movie (*6*). Unweighted full-dose images were further used to estimate the CTF parameters using CTER (*7*) (SPHIRE) (*8*). Dose weighted full-dose images were used for all other steps of image processing. ClpXP1/2 dimers were picked automatically using EMAN2’s (*9*) neuralnet e2boxer. Further data processing was performed using the software package SPHIRE (*10*). After inspection of micrographs using the CTF-assessment-GUI, 273,300 single particles were selected for further processing. The particle stack was subjected to 2D-clustering using ISAC2 (SPHIRE), resulting in a “clean” stack of 143,901 single particles producing stable and reproducible 2D-class averages. The 2D class-averages were used to calculate a 3D volume, using VIPER. After masking, this volume was used as the reference for a 3D refinement using Meridien (SPHIRE), which resulted in a 13 Å density map, as estimated by the “gold-standard” FSC. In agreement to the 2D clustering results (Supplementary Video 1), further 3D clustering using Sort3D (SPHIRE) confirmed that the ClpXP1/2 dimer is a continuously flexible structure (Supplementary Figure 1g). Independent refinement of the resulting subsets did not, however, further improve the resolution of the volume.

We then manually picked the ClpXP1/2 monomers within each ClpXP1/2-dimer for 10 representative micrographs of the dataset and used these data to train crYOLO (*11*), which then automatically selected 613,322 single particles. After 2D and 3D clustering, a final “clean” stack of 383.927 particles was used for further refinement. During the first rounds of the refinement, we applied local symmetrization of the reference after each refinement round, as previously described (*12*, *13*) *i.e.* after each refinement round the density of ClpP was symmetrized using *D7* symmetry, whereas the density of ClpX was scaled in order to put an additional weight on this region during the asymmetric refinement. Finally, both densities (ClpX and ClpP) were combined and the resulting volume was used as a reference for the subsequent refinement iteration. This procedure was performed during the initial rounds in order to obtain global projection parameters. The user function was not applied during the local refinements. This resulted in a density map with an average resolution of 4 Å, where the resolution of the density decreases towards ClpX (Supplementary Figure 2). The average resolution was calculated between two independently refined “half maps” at the 0.143 FSC criterion. The estimated accuracy of rotation and translation search during the last refinement round was estimated to 1.78° and 1.02 pixels, respectively. Local resolution was computed using the “Local Resolution” tool in SPHIRE. 3D clustering into four groups was performed using the RSORT3D tool of SPHIRE. However, according to the ANOVA analysis, the resulting volumes were not reproducible and were therefore not considered for further analysis. 3D Refinement and Clustering focusing on the density of ClpX, after removing the ClpP signal from the dataset, did also not result into further improvement of the ClpX density. The density of ClpP was auto-sharpened locally using phenix.auto_sharpen (*14*) and filtered to its average resolution of 3.9 Å. The ClpX desnity was filtered to an average resolution of 6.5 Å and sharpened with an ad-hoc b-factor of −240 Å^2.^ Angular distribution plots were computed using SPHIRE. Beautified 2D class averages were computed with 3500 members per group.

### Atomic modelling

We built a homology model of ClpX with SWISS-MODEL (*15*) using ADP-bound E. coli ClpX (PDB-ID 3HWS, Chain A) and ATPγS-bound *E. coli* ClpX (PDB-ID 4I81, Chain B). We then used UCSF Chimera (*16*) to fit the structures of ClpX’s homology model and ClpP1/2 (PDBID 4RYF (*2*) into the cryo-EM density. We used the RosettaES protocol (*17*) to build the missing residues 9-16 for each ClpP2 subunit. Residues 1-2 were manually built in Coot (*18*).

With the complete model, we performed several iterative runs of molecular dynamics flexible fitting (MDFF) (*19*) and manual adjustment with Coot, paying particular attention to the fitting of the IGF loops. In the initial run, we applied 6-fold symmetry to ClpX, allowing regions poorly supported by the density to settle into reasonable conformations. This restraint was later removed. For the final iterations, we also included a step of real-space refinement in Phenix (*20*), to decrease the number of Ramachandran outliers and to fit the atomic B-factors. The necessary files for the MDFF runs were set up with VMD (*21*) and all simulations were performed in NAMD (*22*), using the CHARMM 36m force field (*23*) with the implicit solvation model implemented in NAMD.

For the proper modeling of the structure with MDFF, we included all missing regions of the structures, even if their density does not allow full atomic modeling. After refinement, we removed all those from the final model. The quality of this model was assessed in Phenix, using the Molprobity (*24*) and EMRinger scores (*25*) as well as the overall geometry of the structure.

Sequence conservation was analyzed using the ConSurfserver (*26*). Analysis of the channel pathway was performed with ChExVis (*27*). Electron density maps and models were visualized using Chimera (*16*) and Chimera X (*28*).

### Peptidase assay

In this assay, the degradation of a fluorogenic tripeptide was measured, for which ClpX was not required. 99 μL 1 μM ClpP1/2 was incubated in PZ buffer (25 mM HEPES, 200 mM KCl, 5 mM MgCl2, 1 mM DTT, 10% glycerol, pH 7.6) in flat bottom black 96-well plates for 15 min at 30 °C. 1 μL acetylalanyl-homoarginyl-2-aminooctanoyl-7-amino-4-carbamoylmethylcoumarin (Ac-Ala-hArg-2-Aoc-ACC) substrate (10 mM stock in DMSO) was added and the fluorescence was measured (380 nm, 430 nm) with an infinite M200Pro plate reader (Tecan) at 30 °C. Data were recorded in triplicate and two independent experiments were performed. Peptidase activity was determined by linear regression using Microsoft Excel and plots were made with GraphPad Prism 6.

### Protease assay

Protease assays were carried out in flat bottom white 96-well plates in a final volume of 60 μL. (ClpP1/2)_14_ (0.2 μM), ClpX_6_ (0.4 μM) and ATP regeneration mix (4 mM ATP, 16 mM creatine phosphate, 20 U/mL creatine kinase) were pre-incubated for 15 min at 30 °C in PZ buffer. 0.8 μM eGFP-SsrA substrate was added and fluorescence was measured (485 nm, 535 nm) at 30 °C. Data were recorded in triplicate and at least two independent experiments were performed. Protease activity was determined by linear regression using Microsoft Excel and plots were made with GraphPad Prism 6.

### ATPase assay

90 μL 2 μM ClpX in ATPase buffer (100 mM HEPES, 200 mM KCl, 20mM MgCl_2_, 1 mM DTT, 1 mM NADH, 2 mM phosphoenolpyruvate, 50 U/mL lactate dehydrogenase, 50 U/mL pyruvate kinase, 5% glycerol, pH 7.5) was added to a flat bottom transparent 96-well plate and incubated for 15 min at 37 °C. The reaction was started by the addition of 10 μL 200 mM ATP in 100 mM HEPES, pH 7.5. Absorption at 340 nm was measured at 37 °C. Two independent experiments with three replicates each were carried out. ATPase activity was determined by linear regression using Microsoft Excel after subtraction of the background signal (measurement without ClpX), and the plot was made with GraphPad Prism 6.

### Hydrogen/deuterium exchange mass-spectrometry (HDX-MS)

HDX-MS experiments were performed using an ACQUITY UPLC M-class system equipped with automated HDX technology (Waters). HDX kinetics were determined by taking data points at 0, 10, 60, 600, 1800 and 7200 s at 20 °C. At each data point of the kinetic, 3 μL of a solution of 30 μM „free” ClpP1/2 and „free” ClpX were analyzed and compared to the (ClpXP1/2)_2_ complex (1.4 μM). The respective protein solutions were diluted automatically 1:20 into 99.9% D_2_O-containing buffer (25 mM HEPES, 200 mM KCl, 5 mM MgCl_2_, 0.5 mM ATP, 1 mM TCEP, 5% glycerol, pH 7.6). As reference, all samples were analyzed in H_2_O ‒containing buffers. The reaction mixture was quenched by the addition of 1:1 200 mM KH_2_PO_4_, 200 mM Na_2_HPO_4_, pH 2.3 (titrated with HCl) at 1 °C and 50 μL of the resulting sample were subjected to on-column peptic digest on a Waters Enzymate BEH pepsin column 2.1 × 30 mm at 20 °C. Peptides were separated by reverse phase chromatography at 0 °C in trapping mode using a Waters Acquity UPLC C18 1.7 μm Vangard 2.1 × 5 mm pre-column and a Waters Aquity UPLC BEH C18 1.7 μm 1 × 100 mm separation column. For separation, a gradient increasing the acetonitrile concentration stepwise from 5 to 35% in 6 min, from 35 to 40% in 1 min and from 40 to 95% in 1 min was applied and the eluted peptides were analyzed using an in-line Synapt G2-S QTOF HDMS mass spectrometer (Waters). UPLC was performed in protonated solvents (0.1% formic acid), allowing deuterium to be replaced with hydrogen from side chains and amino/carboxyl termini that exchange much faster than backbone amide linkages (*29*). All experiments were performed in duplicate. Deuterium levels were not corrected for back exchange and are therefore reported as relative deuterium levels (*30*). The use of an automated system, i.e. handling all samples at identical conditions, negotiates the need for back exchange correction. MS data were collected over an m/z range of 100-2000. Mass accuracy was ensured by calibration with Glu-fibrino peptide B (Waters) and peptides were identified by triplicates MSE ramping the collision energy from 20-50 V. MS data were analyzed with the PLGS 3.0.3 and DynamX 3.0 software packages and all spectra were checked manually. For each peptide, relative uptake values were determined as follows: relative uptake [%] = deuterium uptake × 100 / maximal uptake. For each amino acid, the average of the relative uptake of all peptides covering the amino acid was calculated. The difference of the relative deuterium uptake between the “free” and “complex” states was calculated for each amino acid. Data were analyzed and visualized using custom MATLAB and python scripts, UCSF Chimera 1.12 and OriginPro 2016.

